# Selective induction by statins of FAM134B-mediated sarcoplasmic reticulum (SR)-phagy degrades the SR calcium pump SERCA1 and contributes to myopathy

**DOI:** 10.1101/2025.11.26.690680

**Authors:** Liyang Ni, Xiaolin Zhao, Lirong Zheng, Makoto Shimizu, Takashi Sasaki, Hidetoshi Sakurai, Ryuichiro Sato, Yoshio Yamauchi

**Author notes:** **Corresponding author**; Yoshio Yamauchi, Ph.D., Department of Applied Biological Chemistry Graduate School of Agricultural and Life Sciences The University of Tokyo 1-1-1 Yayoi, Bunkyo-ku, Tokyo 113-8657, Japan. Graduate School of Humanities and Sciences, Ochanomizu University, Tokyo, Japan (Makoto Shimizu).

## Abstract

Statins, HMG-CoA reductase inhibitors, are the most prescribed medication for lowering low-density lipoprotein cholesterol. Although statin intolerance due to statin-associated muscle symptoms (SAMS) has been a clinically important issue for decades, the molecular mechanisms of statin myopathy remain incompletely understood. To seek to clarify how statins induce myopathy, here we perform transcriptome analysis and identify FAM134B short isoform (FAM134B-S), an endoplasmic reticulum (ER)-phagy receptor, as a statin-inducible gene. FAM134B-S expression is regulated by the mevalonate pathway and sterol regulatory element-binding proteins. We further show that FAM134B plays a critical role in the regulation of skeletal muscle integrity both in the steady-state and statin-treated conditions in human iPS cell-derived myocytes and mice. Mechanistically, FAM134B-S interacts with the sarcoplasmic reticulum (SR) calcium pump SERCA1 and promotes its autophagic degradation. Collectively, our work reveals a hitherto undisclosed mechanism of statin myopathy; blocking the mevalonate pathway induces SERCA1 degradation through FAM134B-mediated SR-phagy, exerting myotoxicity.

## Introduction

Statins, 3-hydroxy-3-methylglutaryl CoA (HMG-CoA) reductase (HMGCR) inhibitors, are the most prescribed drugs for reducing the risk of atherosclerotic cardiovascular diseases by lowering low-density lipoprotein (LDL) cholesterol levels.^1^ Mechanistically, statins activate sterol regulatory element-binding protein (SREBP) and upregulate LDL receptor expression, leading to LDL clearance in the liver.^2^ Although statins are generally well-tolerated and safe for most statin takers, they occasionally exert adverse effects. One of the most common complaints of people taking statins is statin-associated muscle symptoms (SAMSs), ranging from asymptomatic elevations of creatine kinase without muscle pain to myalgia, myositis, myopathy, and the most severe form of statin-induced myopathy (SIM), rhabdomyolysis.^3,4^ In the worst cases, 0.1 percent of patients receiving cerivastatin therapy developed lethal rhabdomyolysis, resulting in the withdrawal of cerivastatin from the global market in 2001.^5^ Furthermore, SAMS is the major background of statin intolerance, which can in turn increase the risk of CVD in patients who cease statin therapy.^3,4^ Therefore, elucidating the molecular mechanisms of SAMS is clinically important for overcoming statin intolerance.

HMGCR is the rate-limiting enzyme of the mevalonate pathway that biosynthesizes sterols and non-sterol isoprenoids, including cholesterol and geranylgeranyl pyrophosphate (GGPP).^6^ SIM suggests that the mevalonate pathway is an important metabolic pathway for maintaining skeletal muscle homeostasis. This hypothesis is supported by genetic evidence from humans and mice;^7–9^ skeletal muscle-specific *Hmgcr* knockout mice exert severe muscle loss and myotoxicity.^7^ In addition, recent studies have discovered the *HMGCR* gene mutations in patients with the late-onset limb-girdle muscular dystrophy (LGMD).^8,9^ The administration of mevalonolactone, the lactone form of the HMGCR enzyme product mevalonate, at least partly restores myopathy in both LGMD patients and skeletal muscle-specific *Hmgcr* knockout mice.^7,8^ These findings highlight the physiological and pathophysiological importance of the mevalonate pathway in skeletal muscle. Studies showed that statins exert multiple myotoxic effects in a mevalonate pathway-dependent manner; they promote proteasomal degradation of muscle proteins through upregulating skeletal muscle-specific E3 ubiquitin ligases, Atrogin-1 (also known as FBOX32 or MAFbx) and MuRF1 (also known as TRIM63), but suppress protein synthesis by inhibiting the Akt-mTORC1 axis, which leads to the disruption of proteostasis, a crucial process for maintaining skeletal muscle mass.^10–14^ A number of studies have been conducted to elucidate the mechanism of statin myopathy, yet the roles of HMGCR and the mevalonate pathway in skeletal muscle remain incompletely understood.

Autophagy is an evolutionally conserved, important metabolic process in response to nutritional and metabolic stresses or stimuli and thus plays critical roles in human health and disease.^15,16^ It degrades and recycles various cellular components, including proteins, lipids, and even cellular organelles.^17,18^ In addition to the ubiquitin-proteasome systems, autophagy is also essential for skeletal muscle homeostasis.^19^ In mice, deletion of ATG7, an essential protein for initiating autophagy, in skeletal muscle causes severe muscle atrophy.^20^ Recent studies have identified mutations in the *ATG7* gene in humans, and these patients also exhibit myopathy.^21^ These findings have disclosed a crucial role of autophagy in skeletal muscle. Several studies have reported that statins induce autophagy in skeletal muscle,^22–24^ but its mechanism and significance remain largely unknown.

Autophagy is classified into non-selective and selective autophagies; non-selective autophagy degrades bulk cytoplasmic materials, whereas selective autophagy engulfs specific cellular compartments such as mitochondria and endoplasmic reticulum (ER) or specific substrate molecules and delivers them to the lysosome for degradation.^25,26^ ER-phagy is a subtype of selective autophagy that degrades ER fragments and proteins.^27,28^ ER-bound proteins play a critical role as ER-phagy receptors in the initiation of ER-phagy. Ten ER-phagy receptors, including FAM134B which is the first receptor discovered^29^, have been identified so far, and these receptors participate in distinct types of ER-phagy^27^. Although molecular mechanisms of ER-phagy have been extensively studied in this decade and are beginning to be delineated, in vivo significances of ER-phagy remain largely unknown in most tissues, including skeletal muscle. Intriguingly, in skeletal muscle, the specific form of ER called sarcoplasmic reticulum (SR) plays an essential role in muscle functions by controlling calcium ion flux and storage.^30^ However, how SR homeostasis is precisely regulated remains poorly clarified.

We recently established an in vitro model of SIM using human iPS cell-derived myocytes (hiPSC-MCs) and demonstrated that statins impair skeletal muscle proteostasis.^10^ In this study, to seek a novel mechanism involved in SIM, we perform RNA-sequencing and bioinformatics analysis using our hiPSC-based model and identify FAM134B that is selectively upregulated by statins in a manner dependent on the mevalonate pathway. Our findings show that FAM134B facilitates the degradation of SERCA1, an SR protein playing an essential role in SR calcium flux, and suggest that the increase in SERCA1 degradation by FAM134B-mediated ER-phagy contributes to statin myopathy.

## Results

### Statins selectively upregulate FAM134B-S expression in hiPSC-MCs

We recently established an hiPSC-based SIM model and showed that statins disrupt proteostasis.^10^ To further explore the mechanism of SIM, we performed RNA-sequencing analyses on two lines of hiPSC-MCs (414C2 and 409B2 lines) treated with cerivastatin in the presence or absence of MVA, the product of HMGCR **(Figures 1 and S1)**. Gene expression profiles showed clear discrimination between control and statin-treated groups, and MVA supplementation to statin-treated cells reversed gene expression profiles, resulting in a similar expression profile to the control group **(Figure 1A)**. Cerivastatin treatment resulted in the change in the expression of 1,051 and 524 genes, which are approximately 6.3% (1,051 out of 16,757 genes) and 3.1% (524 out of 17,012 genes) in hiPSC-MCs-414C2 and hiPSC-MCs-409B2, respectively (**Figure 1B**). On the other hand, adding MVA reversed the expression of half of the genes (**Figure 1B**). These results suggest that statins cause alterations in the expression of a number of genes in an MVA pathway-dependent manner.

**Figure 1.**
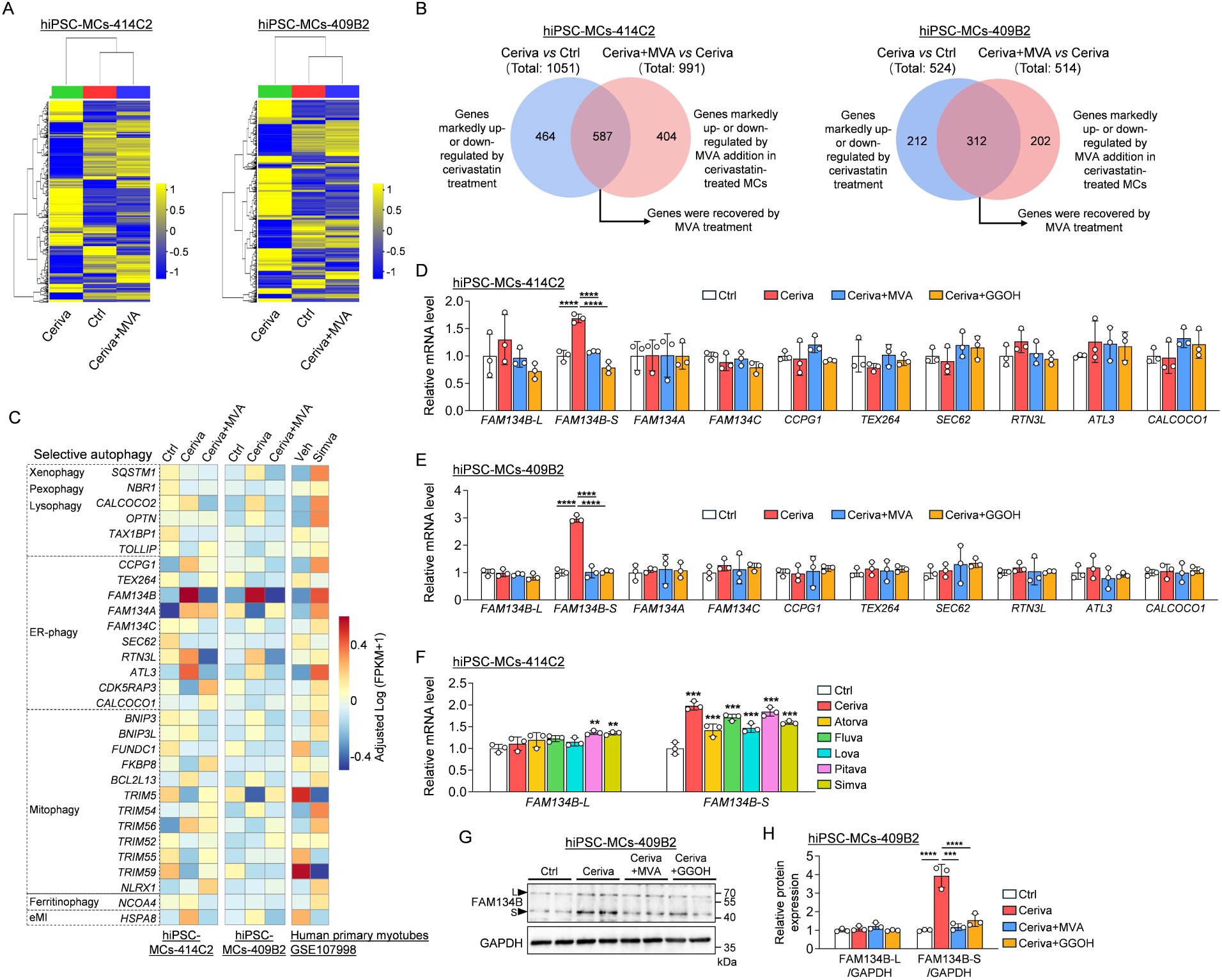
Cerivastatin selectively upregulates FAM134B-S expression in hiPSC-MCs. hiPSC-MCs (414C2 and 409B2) were treated with 5 μM cerivastatin (Ceriva) in the presence or absence of 200 μM MVA or 100 μM geranylgeraniol (GGOH) for 16 h. (A–C) RNA-seq analysis was performed on hiPSC-MCs (414C2 and 409B2) treated with or without cerivastatin and MVA. RNA-seq data are shown as Heatmap (A) and Venn diagrams (B). Genes involved in selective autophagy were selected, and the results are shown as Heatmap (C). Results of human primary myotubes treated with simvastatin (GSE107998) are included. (D, E) mRNA expression of ER-phagy receptors in the hiPSC-MCs-414C2 (D) and -409B2 (E). n=3 per group. (F) hiPSC-MCs-414C2 were treated with different statins (cerivastatin, atorvastatin, fluvastatin, lovastatin, pitavastatin, and simvastatin) at 5 μM for 16 h. mRNA levels of *FAM134B-L* and *FAM134B-S* were analyzed by RT-qPCR. n=3 per group. (G, H) Immunoblot images (G) and quantification data (H) of FAM134B in hiPSC-MCs-409B2. GAPDH was used as an internal control. n=3 per group for quantification. Data shown are mean ± S.D. Statistical analyses were performed by one-way ANOVA with a Dunnett post hoc test. *, p < 0.05, **, p < 0.01, ***, p < 0.001, ****, p < 0.0001 vs cerivastatin-treated group.

To further analyze the transcriptome data, we next performed Kyoto Encyclopedia of Genes and Genomes (KEGG) pathway analysis and found that blocking the MVA pathway by cerivastatin altered various KEGG pathways, including PI3K-Akt signaling pathway, phagosome, mitophagy, mitogen-activated protein kinase (MAPK) pathway, and FOXO signaling pathway (**Figure S1A**). Since most of these pathways are related to autophagy,^31^ and statins disrupt proteostasis in hiPSC-MCs,^10^ we focused on genes related to autophagy, particularly selective autophagy, which maintains tissue homeostasis. Unbiased cluster analysis revealed that among 30 selective autophagy-related genes, the expression of *FAM134B*, which encodes the ER-phagy receptor FAM134B, is selectively upregulated by cerivastatin in both hiPSC-MCs-414C2 and hiPSC-MCs-409B2, and the increase was cancelled by adding MVA (**Figure 1C**). Analyzing a public database (GSE107998) also showed that simvastatin increases *FAM134B* mRNA levels in human primary myotubes, confirming our results with hiPSC-MCs (**Figure 1C**). The human *FAM134B* gene encodes at least two distinct isoforms through the alternative splicing: a full-length 60 kDa (1493 amino acid-length) and an NH₂-terminal truncated 42 kDa (1070 amino acid-length) proteins,^32^ hereafter referred to as FAM134B-long form (FAM134B-L) and FAM134B-short form (FAM134B-S), respectively. To determine which isoforms are upregulated by cerivastatin, we performed qRT-PCR and immunoblotting analysis and found that cerivastatin markedly increases mRNA and protein levels of FAM134B-S in a dose-dependent manner without affecting FAM134B-L expression in hiPSC-MCs (**Figures 1D, E, S1B, C**). Consistent with RNA-seq results, cerivastatin had no significant effects on the mRNA levels of other ER-phagy receptors, including *FAM134A*, *FAM134C*, *CCPG1*, *TEX264*, *SEC62*, *RTN3L*, *ATL3,* and *CALCOCO1* (**Figure 1D, E**), supporting our notion that statins selectively induce FAM134B-S expression. Furthermore, in addition to cerivastatin, atorvastatin, fluvastatin, lovastatin, pitavastatin, and simvastatin also increased mRNA levels of the *FAM134B-S* with only subtle effects on those of *FAM134B-L* in hiPSC-MCs (**Figure 1F**). The increase in FAM134B-S expression by cerivastatin was fully restored by the addition of MVA or GGOH in hiPSC-MCs (**Figures 1D–H, S1D, E**). Similar results were obtained in HeLa cells; the expression of FAM134B-S was increased by cerivastatin, but the upregulation was cancelled by MVA supplementation (**Figure S1F, G**). These results demonstrate that blocking the MVA pathway selectively induces the expression of FAM134B-S, not FAM134B-L or other ER-phagy receptors.

### Statin induces FAM134B-S expression in skeletal muscle in vivo

To investigate whether statins also increase FAM134B-S expression in skeletal muscle in vivo, mice were treated daily with either atorvastatin or fluvastatin (statins used clinically) for 2 weeks (**Figure S2A**). After 7 and 14 days of daily statin administration, mice exhibited significant reductions in grip strength without influencing body weight **(Figure S2B, C)**, establishing statin-induced myopathy in mice. We therefore examined FAM134B-S and -L expression in skeletal muscle (tibialis anterior (TA)) and found that FAM134B-S expression was induced by both atorvastatin and fluvastatin (**Figure S2D, E**). FAM134B-L was nearly undetectable in this tissue, which is consistent with previous studies showing that FAM134B-S is a predominant form of FAM134B in skeletal muscle.^33^ Immunoblot analysis also showed that the expression of Atrogin-1, a skeletal muscle-specific ubiquitin ligase E3, and LC3B-II was also increased by these statins. These results suggest that statins induce not only the ubiquitin-proteasome pathway but also ER-phagy in vivo.

We and others have shown that deficiency of GGPP, not cholesterol, is the major cause of SIM.^10,34^ To investigate whether FAM134B-S expression is regulated by the mevalonate pathway in skeletal muscle, mice were injected with fluvastatin with or without supplementation of mavalonolactone (MLN), a lactone form of MVA, or GGOH for 2 weeks (**Figure 2A**). Fluvastatin alone or co-administration with MLN or GGOH had no effects on body weight (**Figure 2B**). Consistent with **Figure S2C**, fluvastatin reduced grip strength and weight of fast-twitch muscles (quadriceps (Quad), gastrocnemius (Gas) and TA) but not in slow-twitch muscle (soleus (Sol)), and co-administration of fluvastatin with MLN or GGOH partly improved grip strength and muscle losses (**Figure 2C, D**). In addition, fluvastatin markedly decreased myofiber cross-sectional area (CSA), while MLN or GGOH supplementation cancelled the reduction (**Figure 2E, F, G**). Furthermore, mRNA levels of *Fbxo32* (which encodes Atrogin-1) but not *Trim63* (which encodes another skeletal muscle-specific E3 ligase, MuRF1) were upregulated in TA in a manner dependent on the MVA pathway (**Figure 2H**), which is consistent with our previous results in hiPSC-MCs.^10^ All these results support the notion that GGPP deficiency is the primary cause of SIM. We then sought to determine whether statins increase FAM134B-S expression in a mevalonate pathway-dependent manner in skeletal muscle. We examined *FAM134B-S* mRNA levels in Quad, Gas, TA, and Sol, and found that fluvastatin markedly upregulated its mRNA levels in fast-twitch muscle with no significant change in Sol muscle (**Figure 2I**). Immunoblot analysis showed that in addition to LC3B-II and Atrogin-1, fluvastatin increased FAM134B-S expression in TA (**Figure 2J, K**), suggesting both proteasomal and autophagic pathways are activated by statins. The increase in FAM134B expression by fluvastatin in the ER (KDEL-positive organelles) of TA was further confirmed by immunofluorescent staining (**Figure 2L**). The upregulation of FAM134B-S expression in the fast-twitch muscle by fluvastatin was restored by co-treatment with MLN or GGOH (**Figure 2I, J, K**). Taken together, consistent with the observations in hiPSC-MCs, the statin-mediated upregulation of FAM134B-S is dependent on the MVA pathway in vivo.

**Figure 2.**
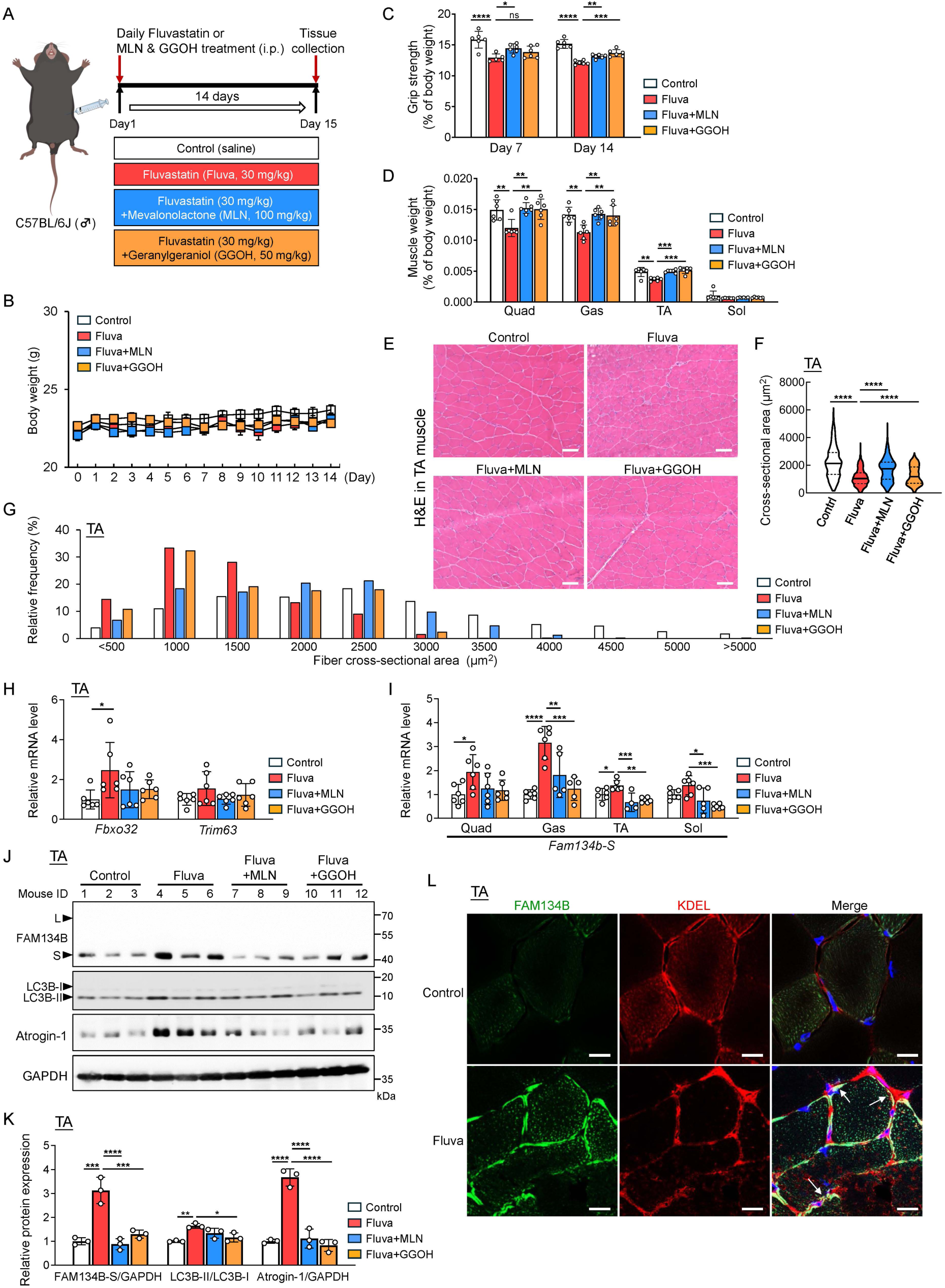
Statin induces FAM134B-S expression in skeletal muscle in vivo. (A) Schematic diagram of experimental design. Mice were treated with or without fluvastatin and MLN or GGOH for 14 days. (B) Body weight was measured daily during the treatments (n=6). (C) Grip strength was measured on days 7 and 14 and normalized to body weight (n=6). (D) Weights of quadriceps (Quad), gastrocnemius (Gas), tibialis anterior (TA), and soleus (Sol) were normalized to body weight (n=6). (E–G) Representative images of H&E staining (E), muscle fiber cross-sectional area (CSA) (F), and distribution of fiber CSA (G) in TA muscle are shown (n=584–1,100 fibers from 3 mice each). Scale bar, 50 μm. (H, I) mRNA levels of *Fbxo32* and *Trim63* in the TA (H) and *Fam134b-S* in the Quad, Gas, TA, and Sol (I) (n=6). (J, K) Protein expression of FAM134B, LC3B, and Atrogin-1 in TA muscles (J). The intensity of bands normalized to GAPDH was quantified (K) (n=3). (L) Expression and localization of FAM134B in TA muscles. Shown are representative confocal images of FAM134B (green) and KDEL-positive ER (red). Data shown as mean ± S.D. Statistical analyses were performed by one-way ANOVA with Dunnett post hoc test. *, p < 0.05, **, p < 0.01, ***, p < 0.001, ****, p < 0.0001 vs Fluvastatin.

### FAM134B-S expression is regulated by SREBPs

The transcription factors SREBPs upregulate genes involved in cholesterol and fatty acid synthesis by binding to the sterol regulatory element (SRE) in the promoter region of target genes.^35^ Since statins activate SREBP-2 processing in hiPSC-MCs,^10^ we asked whether SREBPs regulate FAM134B-S expression. We first cloned -3,000 bp promoter regions of human FMA134B-L and FAM134B-S into a luciferase reporter plasmid and performed the luciferase reporter assay in HEK293T cells (**Figure 3A, S3A**). The results showed that forced expression of either SREBP-1a or SREBP-2 mature form strongly increases the reporter activity for human FAM134B-S promoter, but not human FAM134B-L promoter, suggesting that SREBP-1a and SREBP-2 directly upregulate FAM134B-S expression. As expected, the human *FAM134B-S* promoter region contains nine SRE-like sequences (**Figure S3A**). To confirm the results, we examined whether SREBP-1a and SREBP-2 increase mRNA and protein expression of FAM134B in HeLa cells that express both FAM134B-L and -S endogenously. Overexpression of either SREBP-1a or SREBP-2 mature form selectively induced FAM134B-S expression at both mRNA (**Figure 3B**) and protein levels (**Figure 3C, D**). We recently showed that statins inhibit the protein kinase Akt and activate forkhead box O1 (FOXO1) in hiPSC-MCs.^10^ Therefore, we next tested whether FOXO1 and other FOXO family transcription factors also regulate FAM134B expression. Contrary to our expectation, overexpression of FOXO1, FOXO3a, or FOXO4 increased neither the promoter activity of FAM134B-S and FAM134B-L (**Figure S3B**) nor mRNA levels of both isoforms (**Figure S3C**), indicating that FOXO transcription factors do not directly regulate FAM134B expression. Collectively, our findings suggest that SREBP-1a and SREBP-2 selectively induce FAM134B-S expression.

**Figure 3.**
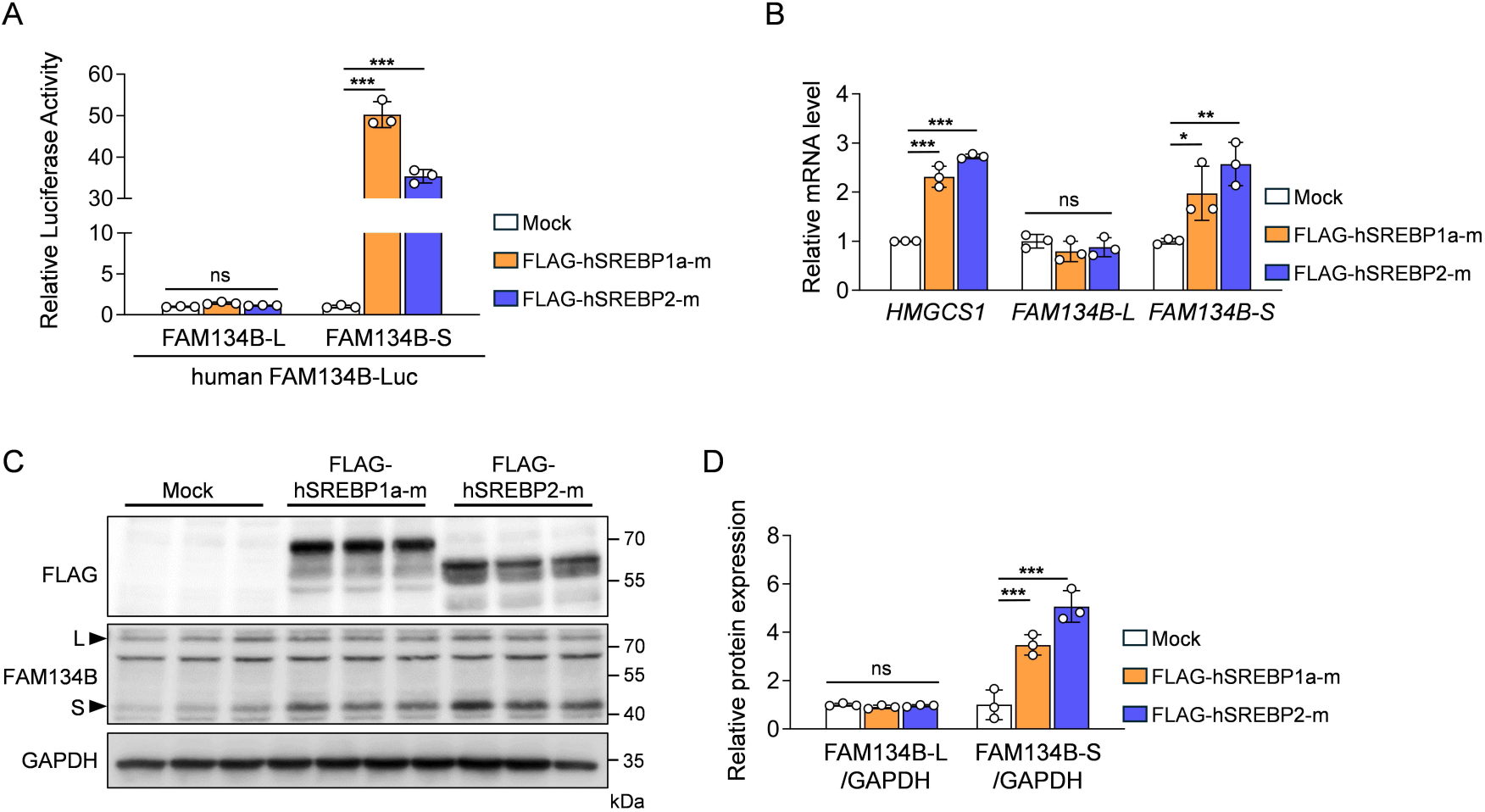
SREBPs upregulate FAM134B-S expression. (A) Luciferase reporter assay was conducted in HEK293T cells. Cells were transfected with human *FAM134B-L* or human *FAM134B-S* promoter gene luciferase reporter plasmid along with an expression vector for either human SREBP-1a or SREBP-2 mature forms. The luciferase activity was measured 24 h after transfection (n=3). (B–D) FLAG-hSREBP1a or FLAG-hSREBP2 mature form was forcedly expressed in HeLa cells. mRNA levels of *HMGCS1*, *FAM134B-L*, and *FAM134B-S* were analyzed by RT-qPCR (n=3) (B). Protein expressions of FLAG and FAM134B were analyzed by immunoblotting (C). The intensity of FAM134B-S bands normalized to GAPDH (D) (n=3). Data shown as mean ± S.D. Statistical analyses were performed by one-way ANOVA with a Dunnett post hoc test. *, p < 0.05, **, p < 0.01, ***, p < 0.001 vs mock.

### Statin promotes FAM134B-mediated ER-phagy

An increase in FAM134B expression itself can induce ER-phagy.^29,36^ We therefore sought to determine whether statin-dependent upregulation of FAM134B-S induces ER-phagy in hiPSC-MCs and other cells. Immunofluorescence staining experiments showed that in response to cerivastatin treatment, massive accumulation of FAM134B in KDEL-positive organelles was observed in the ER and KDEL-positive punctate structures, which were diminished by the addition of MVA or GGOH (**Figure S4A**). To determine whether statins induce autophagy and ER-phagy, we employed DAPGreen, a fluorescent small molecule that can visualize autophagosomes and autolysosomes.^37^ First, we examined the responses of DAPGreen to starvation or cerivastatin in RC-13 (rhabdomyosarcoma cells) **(Figure S4B)** and HeLa cells **(Figure S4C)**. As expected, starvation markedly induced DAPGreen signals in these cells, confirming its usefulness. We next examined the effect of cerivastatin on DAPGreen and found that cerivastatin progressively increased DAPGreen signals in RC-13 and HeLa cells (**Figure S4B, C**), indicating the induction of autophagy by cerivastatin. Similar results were also observed in hiPSC-MCs; in addition to starvation, cerivastatin treatment increased DAPGreen signals in a time-dependent manner, and the increases were restored by the addition of MVA (**Figure S4D, E**), suggesting that statin-induced autophagy depends on the mevalonate pathway. Further analysis showed that prolonged cerivastatin treatment induces colocalization of DAPGreen signals with LysoTracker, a lysosome marker (**Figure S4D**), indicating the formation of the autolysosome.

Given that FAM134B is an ER-phagy receptor, we next asked whether cerivastatin induces FAM134B-dependent ER-phagy in hiPSC-MCs. Incubation of hiPSC-MCs with cerivastatin for 16 h markedly increased both DAPGreen and FAM134B signals (**Figure 4A**). We also found the marked increase in puncta that contain both FAM134B and DAPGreen, suggesting that cerivastatin upregulates FAM134B-mediated ER-phagy. The increases in DAPGreen and FAM134B signals were abrogated by supplementing MVA or GGOH, which indicates cerivastatin induces FAM134B-mediated ER-phagy via the depletion of GGPP. To further corroborate these results, we performed transmission electron microscopy (TEM) analysis in hiPSC-MCs (**Figure 4B, S4F**). The cerivastatin treatment resulted in the accumulation of autolysosome-like structures that contain cellular components, including those resembling the ribosome-attached rough ER, and such structures were not observed in myocytes co-treated with cerivastatin and MVA or GGOH. Collectively, we conclude that statins induce FAM134B-mediated ER-phagy in human myocytes in a manner dependent on the mevalonate pathway.

**Figure 4.**
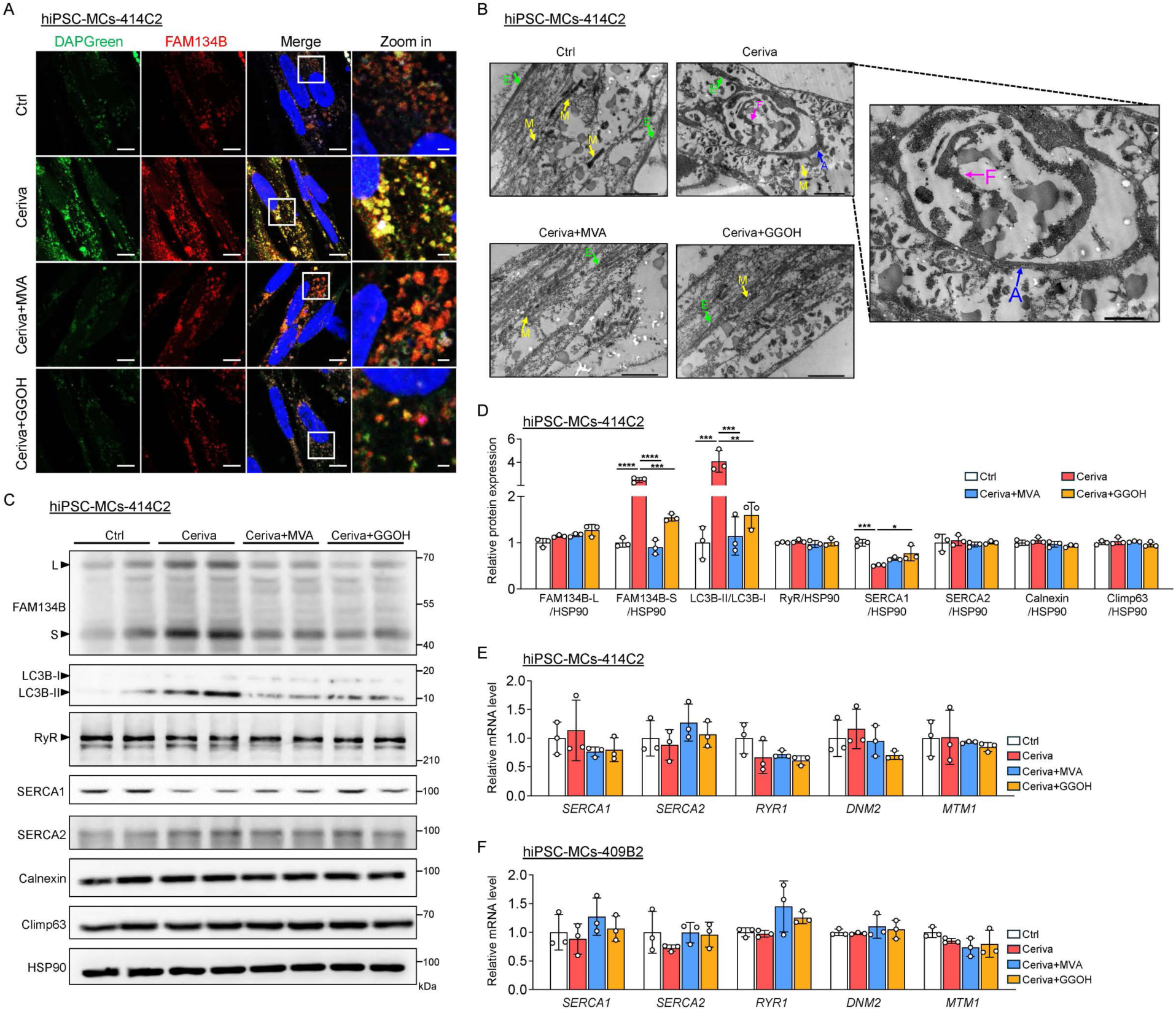
Statin promotes FAM134B-mediated ER-phagy. (A) Localization of DAPGreen (green) and FAM134B (red) in hiPSC-MCs-414C2 treated with 5 μM cerivastatin in the presence or absence of MVA (200 μM) or GGOH (100 μM) for 16 h. Scale bar, 5 μm. (B) TEM images of hiPSC-MCs-414C2 treated as above. Scale bar, 1 μm. E, ER; M, mitochondria; A, autolysosome; F, ER-like fragments. (C, D) Protein expression of FAM134B, LC3B, RyR, SERCA1, SERCA2, Calnexin and Climp63 (C) in hiPSC-MCs-414C2 treated as above. HSP90 was used as an internal control. Quantification data are shown in D (n=3). (E, F) mRNA expression of SR-related genes (*SERCA1*, *SERCA2*, *RYR1*, *DNM2,* and *MTM1*) in the hiPSC-MCs-414C2 (E) and hiPSC-MCs-409B2 (F) treated with cerivastatin, MVA or GGOH as above (n=3). Data shown as mean ± S.D. Statistical analyses were performed by one-way ANOVA with Dunnett post hoc test. **, p < 0.01, ***, p < 0.001, ****, p < 0.0001 vs Cerivastatin.

### Statin induces SERCA1 degradation through FAM134B-mediated SR-phagy

In skeletal muscle, a specialized ER form known as the sarcoplasmic reticulum (SR) is dedicated to maintaining calcium homeostasis.^30^ Based on the above results, we wondered whether statins induce autophagic degradation of ER/SR proteins crucial for skeletal muscle functions. To test this hypothesis, we assessed protein and mRNA levels of ER/SR proteins, including sarcoplasmic/endoplasmic reticulum calcium ATPase 1 (SERCA1), SERCA2, ryanodine receptor (RyR), Calnexin and Climp63. The treatment of hiPSC-MCs with cerivastatin increased FAM134B-S in a mevalonate pathway-dependent manner (**Figure 4C–F**). On the other hand, cerivastatin significantly reduced the protein expression of SERCA1, a protein highly expressed in fast-twitch muscle, without affecting its mRNA levels, and this reduction was restored by the addition of GGOH. Other ER/SR resident proteins tested were not affected by cerivastatin both at protein and mRNA levels. These results suggest that SERCA1 protein contents are selectively decreased by statins in a manner dependent on the MVA pathway.

We next sought to determine whether cerivastatin induces the degradation of SERCA1. The degradation rate of the ER/SR proteins was first examined in hiPSC-MCs using the translation inhibitor cycloheximide (CHX). We found that SERCA1 is a protein with a shorter half-life than SERCA2; the CHX treatment for 1 h resulted in the reduction of SERCA1 protein by 60% (**Figure S5A, B**). The results also showed that FAM134B-S and other ER/SR proteins also decreased by more than 50% within 3-h CHX treatment. Next, we examined whether cerivastatin induces SERCA1 degradation via autophagy. **Figures 5A and B** show that the reduction in SERCA1 protein by cerivastatin was inhibited by Bafilomycin A1, a lysosome/autolysosome inhibitor, suggesting that cerivastatin promotes the degradation of SERCA1 through autophagy. FAM134B-S degradation was also attenuated by Bafilomycin A1 as reported.^33^ Bafilomycin A1 did not affect SERCA1 abundance in the absence of cerivastatin although it increased FAM134B-S, presumably because SERCA1 is also degraded by the proteasome pathway.^38^ Other ER/SR proteins tested were unchanged by Bafilomycin A1, suggesting that these proteins are not degraded by autophagy. To determine the autophagic degradation of SERCA1, we generated an ER-phagy reporter protein by fusing SERCA1 with tandem mCherry and eGFP (mCherry–eGFP–SERCA1) **(Figure 5C)**, which was validated by immunoblotting in C2C12 myoblasts **(Figure S6A)**. Under low ER-phagy flux, both eGFP and mCherry emit fluorescence, appearing as yellow reticular signals in the ER. In contrast, high ER-phagy promotes the delivery of ER proteins to the lysosomes, where acid-labile GFP fluorescence is rapidly quenched, resulting in an increase in acid-stable mCherry signals in autolysosomes.^29^ Cerivastatin treatment markedly increased the number of mCherry-positive/eGFP-negative red puncta in C2C12 myoblasts, while these red puncta disappeared in cerivastatin-treated cells on the presence of Bafilomycin A1 **(Figure 5D, S6B)**. These results suggest that cerivastatin promotes the delivery of SERCA1 from SR into the lysosomes. Next, we overexpressed FLAG-SERCA1 in C2C12 myotubes and examined its colocalization with the autophagosome/autolysosome marker DAPGreen. The results showed that cerivastatin treatment increases DAPGreen-positive puncta that contain FLAG-SERCA1 (**Figure 5E**), suggesting that SERCA1 is a substrate of SR-phagy. Collectively, these findings demonstrate that cerivastatin induces autophagic degradation of SERCA1.

**Figure 5.**
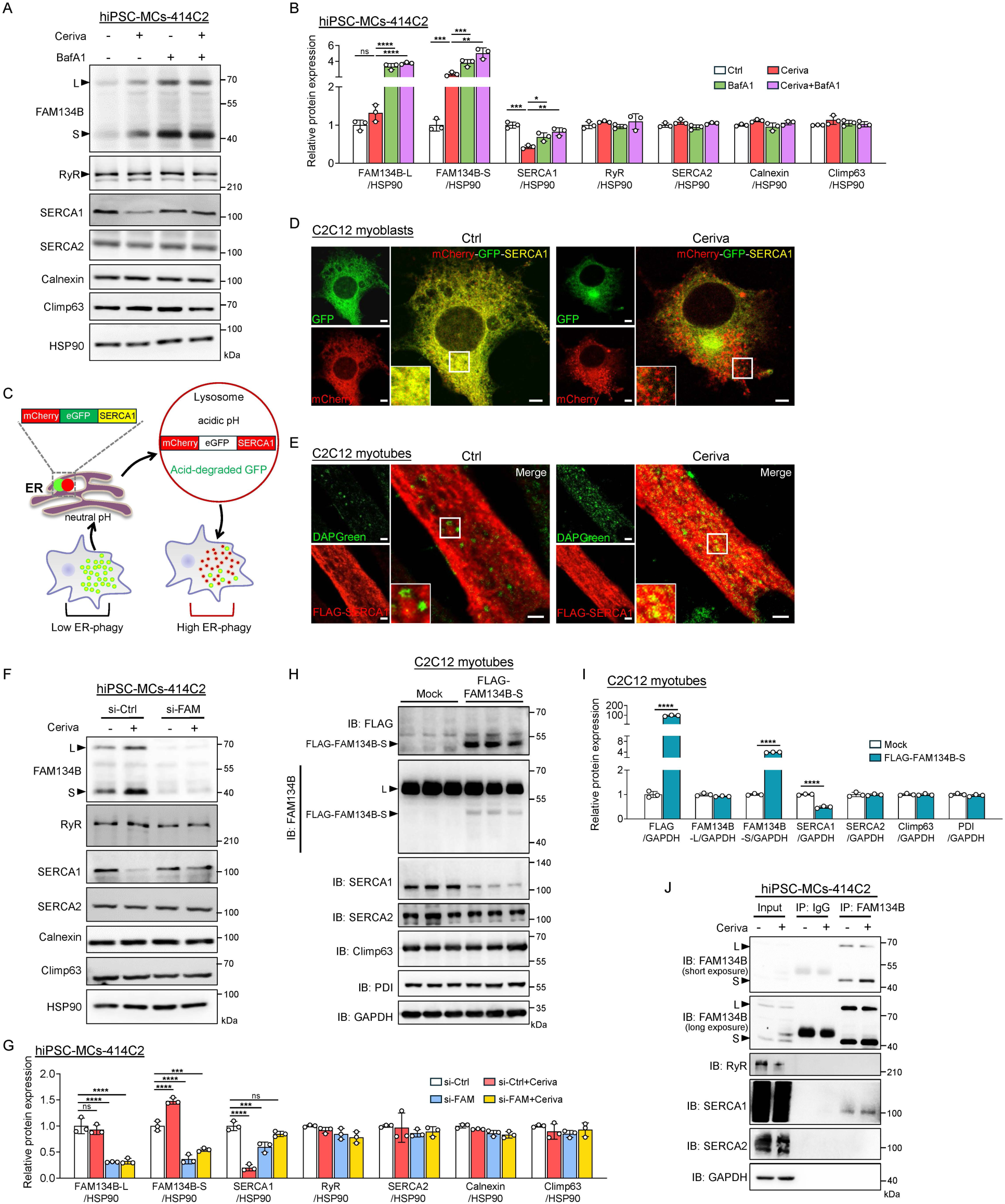
Statin induces SERCA1 degradation through FAM134B-mediated SR-phagy. (A, B) Protein expressions of FAM134B, RyR, SERCA1, SERCA2, Calnexin, and Climp63 in hiPSC-MCs-414C2 treated with 5 μM cerivastatin for 16 h (A). Baf A1 (100 nM) was added 3 h before harvest. HSP90 was used as an internal control, and quantification results are shown in B (n=3). (C) Schematic representation of the ER-phagy reporter mCherry-eGFP-SERCA1. Under steady state, mCherry-eGFP-SERCA1 emits both green and red fluorescence in neutral pH environment. When ER-phagy is activated, the mCherry-eGFP-SERCA1 reporter is transported to acidic autolysosomes and GFP signal is quenched in the acidic condition; the reporter only emits red fluorescence. (D) mCherry-eGFP-SERCA1 was forcedly expressed in C2C12 myoblasts, and cells were treated with 5 μM cerivastatin for 16 h. Representative confocal images of GFP (green) and mCherry (red), scale bar, 5 μm. (E) FLAG-SERCA1 was overexpressed in C2C12 myotubes. The myotubes were incubated with 0.1 μM DAPGreen for 30 min and then treated with or without 5 μM cerivastatin for 16 h. Representative confocal images of DAPGreen (green) and FLAG-SERCA1 (red). scale bar: 5 μm. (F, G) hiPSC-MCs-414C2 were transfected with 25 nM control or FAM134B siRNAs. Cells were treated with or without 5 μM cerivastatin for 16 h. Protein expressions of FAM134B, RyR, SERCA1, SERCA2, Calnexin and Climp63 (F) and quantification data (G) in hiPSC-MCs-414C2. HSP90 was used as an internal control. n=3. (H, I) Protein expressions of FLAG, FAM134B, SERCA1, SERCA2, Climp63, and PDI in C2C12 myotubes overexpressed FLAG-FAM134B-S (H). GAPDH was used as an internal control and the expression of indicated proteins was quantified (I) (n=3). (J) Interaction of FAM134B and SERCA1 in hiPSC-MCs. FAM134B immunoprecipitation was performed as described in Methods. FAM134B, RyR, SERCA1, and SERCA2 were detected by immunoblot. GAPDH was used as an internal control for the inputs. Data shown as mean ± S.D. Statistical analyses were performed by student t-test (I) or one-way ANOVA with Tukey-Kramer post hoc test (B, G). *, p < 0.05, **, p < 0.01, ***, p < 0.001, ****, p < 0.0001.

To investigate the involvement of FAM134B in the autophagic degradation of SERCA1, we silenced FAM134B expression with siRNAs and assessed SERCA1 protein expression in hiPSC-MCs. The results showed that silencing FAM134B selectively protects SERCA1 from cerivastatin-induced degradation (**Figure 5F, G**). FAM134 knockdown did not affect mRNA levels of most ER/SR proteins, including SERCA1 (**Figure S6C**). To determine autophagic degradation of SERCA1 by two FAM134B isoforms, we forcedly expressed either FLAG-FAM134B-S or FAM134B-L-FLAG in C2C12 myotubes **(Figure 5H, I)** and myoblasts **(Figures S6D, E)**, and examined the expression of SERCA1 and other ER/SR proteins. The results show that FAM134B-S overexpression itself markedly reduces SERCA1 expression without affecting other protein contents, suggesting that FAM134B-S induces autophagic degradation of SERCA1. To further confirm this hypothesis, mCherry-eGFP-SERCA1 and FLAG-FAM134B-S were expressed in C2C12 myoblasts, and their localization was examined under a confocal microscopy. The results showed that forced expression of FLAG-FAM134B-S was sufficient to increase mCherry-positive/GFP-negative SERCA1 signals that colocalize with FLAG-FAM134B-S (magenta), while mCherry-positive/GFP-negative red puncta were not observed in BafA1-tretaed myoblasts **(Figure S6F)**. To examine a mechanism by which FAM134B-S selectively induces SERCA1 degradation, we asked whether FAM134B interacts with SERCA1. The immunoprecipitation experiment using anti-FAM134B showed co-immunoprecipitation of endogenous SERCA1 with endogenous FAM134B in hiPSC-MCs (**Figure 5J**). Similar results were obtained in HEK293T cells overexpressing FLAG-SERCA1 and mCherry-FAM134B-S; FLAG-SERCA1 co-immunoprecipitated with mCherry-FAM134B-S (**Figure S6G**). Taken together, these results indicate that cerivastatin induces FAM134B-S-mediated autophagic degradation of SERCA1. Furthermore, since FAM134B is an ER-phagy receptor and SERCA1 is a SR protein expressed in fast-twitch skeletal muscle, we propose to term this myocyte-specific ER-phagy regulating SERCA1 degradation SR-phagy hereafter.

### FAM134B is essential for maintaining the integrity of skeletal muscle

The physiological significance of FAM134B in skeletal muscle remains largely unknown. We first examined the effect of FAM134B silencing in hiPSC-MCs. The knockdown efficiency was greater than 70% both at mRNA and protein levels (**Figure 6A, B**). hiPSC-MCs silenced FAM134B or not were treated with cerivastatin for 16 h, and the diameter of myosin heavy chain (MHC)-positive myocytes was measured. Consistent with our previous report,^10^ cerivastatin reduced the diameter of myocytes (**Figure 6C**). In addition, FAM134B knockdown itself markedly induced atrophy and further aggravated cerivastatin-dependent atrophy. These results suggest that FAM134B plays important roles in maintaining muscle mass at least in vitro.

**Figure 6.**
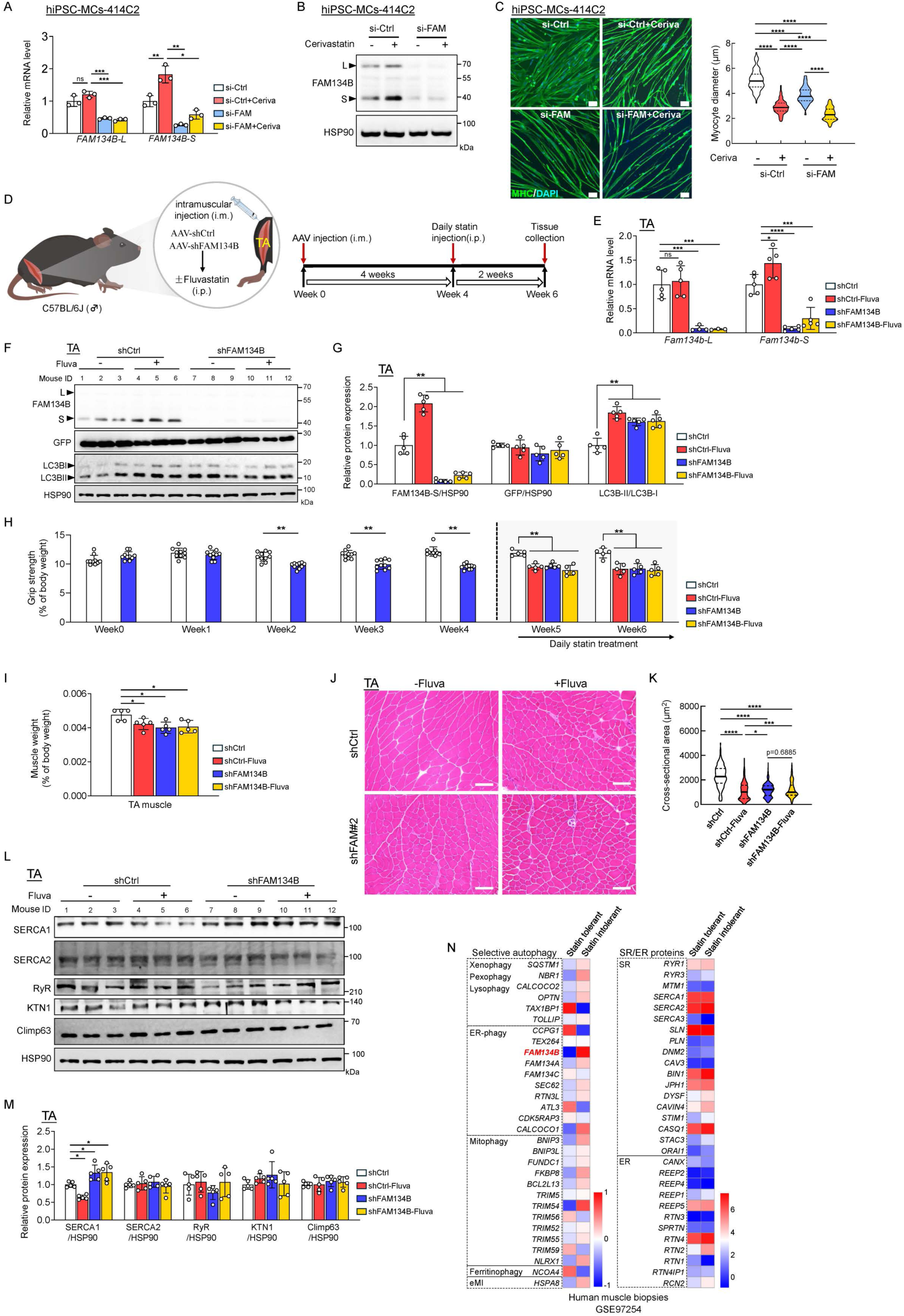
FAM134B is essential for maintaining the integrity of skeletal muscle. (A, B) hiPSC-MCs-414C2 were transfected with control or FAM134B siRNAs, and treated with or without 5 μM cerivastatin for 16 h. mRNA (A) and protein (B) levels of FAM134B were analyzed. HSP90 was used as an internal control (B). n=3 for A. (C) Representative confocal images and myocyte diameters of MHC-positive myocytes. Myocytes were transfected with control or FAM134B siRNAs and treated with cerivastatin as above. Results are expressed in violin plots (n=100). Green, MHC; Blue, DAPI. Scale bar, 50 μm. (D) Schematic of AAV-mediated FAM134B knockdown in the skeletal muscle in mice. (E) mRNA expression of *Fam134b-L* and *Fam134b-S* in the TA from mice infected with either shControl- or shFAM134B-AAVs. Mice were treated or not with fluvastatin for 2 weeks. n=5 per group. (F, G) Immunoblot images (F; 3 mice/treatment) and quantification data (G; n=5) of FAM134B, GFP, and LC3B in the TA. HSP90 was used as an internal control. (H) The grip strength was recorded weekly and normalized to body weight. (I) Weights of TA normalized to body weight are shown (n=5). (J, K) H&E staining images (J) and myofiber CSA (K) in TA. n=485–1101 myofibers from 3 mice each. Scale bar, 50 μm. (L, M) Immunoblot images (L; 3 mice/treatment) and quantification data (M; n=5) of SERCA1, SERCA2, RyR, KTN1, Climp63 in the TA. HSP90 was used as an internal control. (N) Heatmap analysis of mRNA levels of selective autophagy-associated or SR/ER genes in statin-tolerated and -intolerated human muscle biopsies. Data are obtained from the database GSE97254. Data shown as mean ± S.D. Statistical analyses were performed by one-way ANOVA with Tukey-Kramer post hoc test. *, p < 0.05, **, p < 0.01, ***, p < 0.001, ****, p < 0.0001.

We next sought to determine in vivo significance of FAM134B in adult muscle of mice. To specifically silence FAM134B in adult muscle, we took advantage of intramuscular injection of adeno-associated virus (AAV) harboring FAM134B shRNA (shFAM#1 and shFAM#2) into TA (**Figures 6D, S7A**). The successful injection could be confirmed with GFP signals in the hind limb (**Figure S7A**). Both shFAM#1 and #2 markedly reduced the expression of FAM134B-S in TA at both mRNA and protein levels by approximately 80% or more without affecting the expression of *Lc3a, Lc3b*,*Ccpg1*, *Atl3*, and *Tex264,* other ER-phagy receptors (**Figures 6E–G, S7B–E, S8E**). In mouse muscle tissues, FAM134B-L protein was undetectable as reported.^33^ FAM134B knockdown increased LC3B-II in TA, suggesting the induction of autophagy (**Figures 6F, G, S7C, D**). Silencing muscular FAM134B reduced the grip strength of hind limbs from 2 weeks after AAV injection without affecting body weight (**Figures 6H, S7F, G, S8A**). TA mass and cross-sectional area of TA fibers were also markedly decreased by FAM134B knockdown (**Figures 6I–K, S7H–L, S8B–D)**.

We next examined whether statin treatment further affects muscle mass and strength of muscle with FAM134B knockdown. As expected, fluvastatin administration increased FAM134B-S expression, and FAM134B knockdown cancelled the increase in TA (**Figures 6E–G, S7B–D**). In these conditions, FAM134B knockdown did not further decrease grip strength, TA mass, and fiber size in mice treated with fluvastatin (**Figures 6H–K, S7G–L, S8B–D)**, presumably because FAM134B knockdown efficiency was high enough to markedly reduce these indices.

We finally assessed the effect of FAM134B silencing on SERCA1 expression in mouse muscle. Consistent with in vitro data (**Figure 5F, G, S6C**), FAM134B knockdown cancelled statin-dependent reduction of SERCA1 protein expression or even increased its protein levels with no effects on its mRNA levels (**Figures 6L, M, S7M–O, S8E**), suggesting that SERCA1 is a substrate of FAM134B-S-mediated SR-phagy in vivo. To support our findings in hiPSC-MCs and mice, we re-analyzed gene expression profiling data from patients with a background of SIM who underwent rechallenge with statin treatment (GSE97254).^39^ Compared to statin-tolerant participants, patients with statin intolerance who reported muscle pain, cramping, or myalgia following statin re-administration exhibited higher mRNA levels of *FAM134B* without changes in the mRNA expression of most ER/SR-related proteins **(Figure 6N)**. Altogether, our results suggest that statins induce FAM134B-mediated SR-phagy and promote the degradation of SERCA1, which participates in the manifestation of SAMS. Moreover, it is suggested that FAM134B, particularly FAM134B-S solely expressed in skeletal muscle, is a critical autophagy regulator that maintains skeletal muscle homeostasis in adult muscle.

## Discussion

How the mevalonate pathway contributes to muscle integrity remains incompletely understood. Proteostasis is critical for regulating skeletal muscle mass. Using our hiPSC-based statin myopathy model, we recently showed that statins impair proteostasis by disrupting the balance between *de novo* protein synthesis and proteasomal muscle protein degradation.^10^ In this work, to further explore the role of the mevalonate pathway in skeletal muscle and better understand SAMS, we performed transcriptome analysis using our hiPSC model and identified the ER-phagy receptor FAM134B-S as a novel statin-inducible, mevalonate pathway-regulated gene that participates in proteostasis. It is of interest that FAM134B expression in skeletal muscle is upregulated not only in SIM but also in other settings, including cancer cachexia, spinal and bulbar muscular atrophy, fasting, and spaceflight (**Figure 7A)**. Our findings showed that FAM134B-S mediates the degradation of the SR calcium pump SERCA1 and serves as a key regulator of skeletal muscle homeostasis (**Figure 7B**).

**Figure 7.**
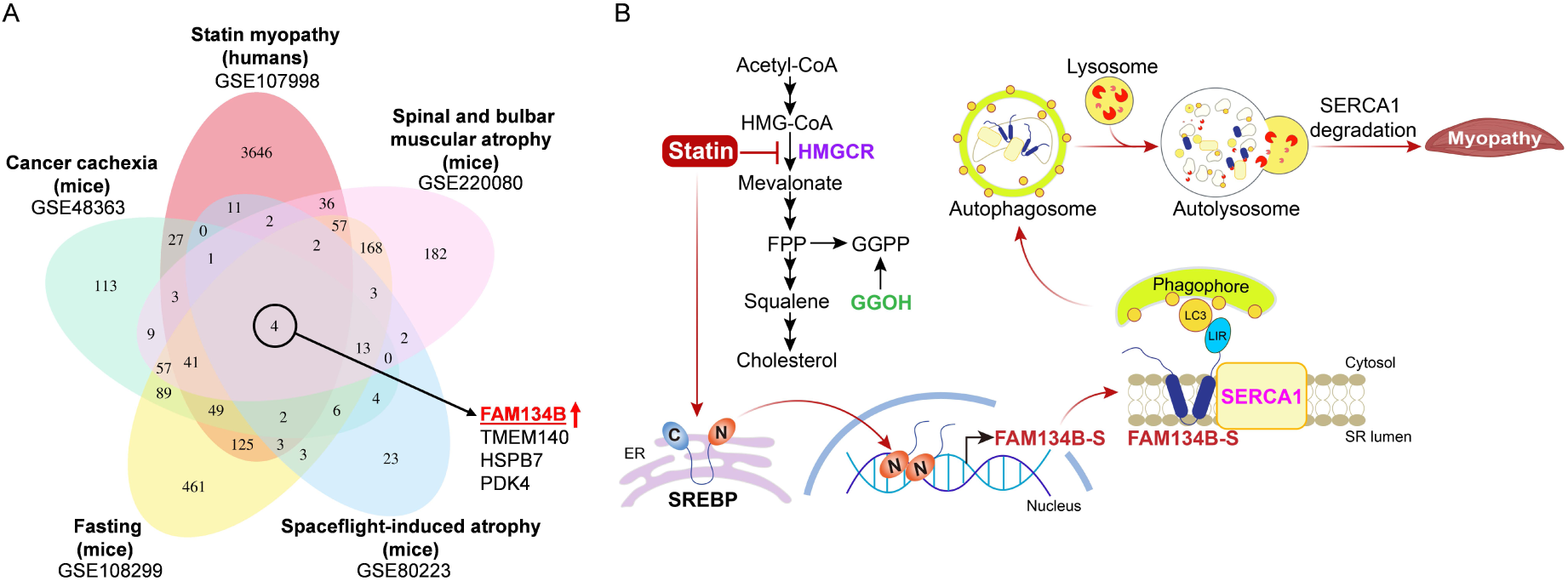
Pathophysiological significance of FAM134B in muscle atrophy. (A) Venn diagram showing the distinct and overlapping genes whose expression is changed in skeletal muscle by the five muscle atrophic conditions in humans and mice. The expression of the four genes, including FAM134B, is commonly altered in these atrophic conditions. (B) Scheme illustrating FAM134B-mediated SR-phagy in statin myopathy. The blockade of the MVA pathway by statins activates SREBP processing and upregulates the FAM134B-S in the skeletal muscle. FAM134B-S induces the SR-phagic degradation of SERCA1, leading to SIM.

In humans and mice, deficiency of ATG7, an essential factor for both non-selective and selective autophagy, leads to myopathy,^20,21^ demonstrating that autophagy plays a crucial role in the regulation of skeletal muscle mass. However, autophagy may have yin-and-yang effects on skeletal muscle. For instance, autophagy is activated and promotes muscle wasting in cancer cachexia, while in aged muscle, decreased autophagic flux results in the accumulation of misfolded proteins and damaged organelles, contributing to sarcopenia.^19^ Moreover, exercise-induced autophagy exerts beneficial effects on skeletal muscle.^40^ Although several studies have shown that statins induce autophagy in several rodent models,^22–24^ little is known about how they induce autophagy in skeletal muscle. Intriguingly, in muscle-specific *Atg7*^-/-^ mice, damaged or aberrant organelles, including swollen SR, are accumulated in skeletal muscle,^20^ implying that autophagy contributes to SR remodeling and clearance. We therefore focused on selective autophagy and sought selective autophagy receptors for which expression is modulated by the mevalonate pathway. Based on our RNA-seq analysis with hiPSC-MCs, we found that the ER-phagy receptor FAM134B-S is selectively induced by statins and that its expression is transcriptionally upregulated by SREBPs. FAM134B contains two functionally important domains; the LC3-interacting region (LIR) at the C-terminus recruits the autophagy machinery to the ER, while the reticulon homology domain (RHD) plays a key role in the ER membrane shaping.^41^ FAM134B is expressed as the full-length long and the N-terminus-truncated short isoforms (FAM134B-L and -S, respectively) through alternative splicing,^32^ and both isoforms are functional as ER-phagy receptor.^36,42^ The expression of the two isoforms is highly cell- and tissue-dependent; FAM134B-L is primarily expressed in the brain, testis, spleen, and prostate, whereas FAM134B-S is restricted in heart, liver, and skeletal muscle in mice.^33^ We showed that FAM134B-S is the major isoform in hiPSC-MCs, suggesting that it is also a predominant isoform in human skeletal muscle. We found that statins upregulate FAM134B-S expression in vitro and in vivo. Since FAM134B expression can drive ER-phagy through ER membrane remodeling by clustering the receptor and an associated protein,^29,36,43,44^ it is conceivable that statin-dependent increase in FAM134B-S promotes SR membrane remodeling for facilitating SR-phagy in skeletal muscle.

Our results also suggest that FAM134B-S expression is regulated by SREBP-1 and SREBP-2, which are activated by statin treatment. We and others have shown that the expression of Atrogin-1 and MuRF1, muscle-specific ubiquitin E3 ligases that induce muscle atrophy and statin myopathy, is regulated by FOXO family transcription factors.^10,45^ Unexpectedly, FOXOs did not upregulate FAM134B expression. On the other hand, FOXOs increase the expression of the essential autophagy gene LC3 by starvation in skeletal muscle.^46^ Based on our results, it is suggested that at least in the setting of statin myopathy, SREBPs play a role in the upregulation of FAM134B-S expression in this tissue. As blocking HMGCR activates FOXOs and SREBPs, we hypothesize that these transcription factors control distinct atrophic genes and convergently induce muscle atrophy in SIM.

What is the pathophysiological significance of SR-phagy induction by statins? Although the physiological and pathophysiological significance of ER-phagy in vivo remains poorly understood, recent studies have shown that FAM134B-mediated ER-phagy remodels the ER during neurogenesis and myogenesis in vitro.^47,48^ We showed that silencing FAM134B expression inhibits autophagic degradation of the muscle-specific SR calcium pump SERCA1, whereas statin-induced expression or forced expression of FAM134B-S promotes the degradation of this pump in a bafilomycin A1-sensitive manner. Thus, these results are consistent with the conclusion that FAM134B-S induces autophagic degradation of SERCA1 in skeletal muscle. Here, we propose to term this skeletal muscle-specific autophagic SR protein degradation pathway “SR-phagy”. Among SR proteins we tested, FAM134B-S enhanced the degradation of SERCA1 only; neither SERCA2 nor other SR proteins tested were unchanged by FAM134B knockdown or overexpression. SERCA1 is a fast-twitch muscle-specific calcium pump, while SERCA2 is predominantly expressed in slow-twitch muscle and cardiac muscle.^49^ The *SERCA1* gene mutations cause Brody myopathy, an inherited muscular disease, and the ablation of SERCA1 in mice shares similar phenotypes with these patients.^50,51^ Although SERCA1 overexpression mitigates phenotypes of muscle diseases in several mouse models, including muscular dystrophy and amyotrophic lateral sclerosis, its overexpression in wild-type mice leads to myopathy.^52–54^ The evidence suggests that fine-tuning the balance of SERCA1 levels and calcium homeostasis plays critical roles in maintaining muscle integrity. Importantly, statins often affect the fast-twitch muscle in humans and rodents and increase cytosolic Ca²⁺ levels due to the disruption of Ca²⁺ storage in the SR.^4,55–59^ These data are consistent with our results that statins reduce the abundance of SERCA1, not SERCA2. All these current and published data support our hypothesis that blocking HMGCR upregulates FAM134B-S, promotes SERCA1 degradation through SR-phagy, and affects fast-twitch muscle. Although the mechanism of LGMD caused by *HMGCR* mutations remains essentially unknown, it is plausible that these patients also exhibit reduced SERCA1 protein levels and impaired calcium homeostasis.

Mutations in the *FAM134B* gene lead to hereditary sensory and autonomic neuropathy type II (HSAN-II), a rare autosomal recessive disorder that affects peripheral sensory and autonomic neurons,^60^ and *FAM134B*^-/-^ mice also show similar phenotypes in the peripheral nerve.^61^ Although these mice exhibit muscle atrophy,^62^ whether this atrophic phenotype is due to peripheral neuropathy or a direct involvement of FAM134B in skeletal muscle homeostasis remains essentially unknown. To exclude the possibility that FAM134B deletion indirectly induces muscle atrophy by peripheral neuropathy, we directly delivered AAV to skeletal muscle to knock down FAM134B in adult skeletal muscle in this work. The results suggest that muscular FAM134B directly participates in myopathy. Thus, it is of interest to test whether patients with HSAN-II also display impairment in SR-phagy and calcium homeostasis in skeletal muscle.

In conclusion, our study identified FAM134B-S as a statin-inducible, mevalonate pathway-regulated gene and demonstrated its key role in skeletal muscle homeostasis. In response to statin treatment, FAM134B-S induces SERCA1 degradation through SR-phagy, a skeletal muscle-specific ER-phagic degradation process. Based on our findings, we conclude that the selective induction of FAM134B-S-mediated SR-phagy participates in the development of SAMS.

## Limitations of this study

Our findings show that statins promote the degradation of the skeletal muscle-specific calcium pump SERCA1 by activating FAM134B-S-mediated SR-phagy, which at least partly participates in the development of SIM. Here, as limitations of this work, we list several outstanding questions to further understand the mechanism and significance of FAM134B-mediated SR-phagy. Firstly, this work does not elucidate a detailed mechanism by which FAM134B-S selectively recognizes and targets SERCA1 for autophagic degradation; understanding the mechanism of SERCA1 degradation will help develop strategies for treating muscular diseases where SR calcium homeostasis is impaired. Moreover, whether muscle atrophy by silencing FAM134B is solely attributed to impaired SERCA1 degradation through SR-phagy remains to be determined. Given the crucial role of SERCA1 in regulating ER calcium flux, its degradation via FAM134B-mediated SR-phagy may also disrupt calcium homeostasis in skeletal muscle—an aspect that was not addressed in this study. It is also important to clarify whether FAM134B-S mediates the autophagic degradation of other SR proteins other than SERCA1.

## Data availability

Data will be made available upon reasonable request.

## Supporting information

This article contains supporting information.

## Acknowledgments

We thank Drs. Atsushi Miyajima and Tohru Itoh for their generous provision of AAV vectors and protocols, and Dr. Satoshi Kimura for his excellent technical assistance with electron microscopy. L.N. was supported by the Otsuka Toshimi Scholarship Foundation.

## Authors contributions

Conceptualization, Y.Y.; Investigation, L.N., X.Z., L.Z., Y.Y.; Formal analysis, L.N., X.Z., Y.Y.; Visualization, L.N., X.Z., Y.Y.; Methodology, L.N., X.Z., L.Z., M.S., T.S., R.S., Y.Y.; Resources, L.N., M.S., H.S., R.S., Y.Y., Writing – original draft, L.N., Y.Y.; Writing – review & editing, L.N., Y.Y.; Funding acquisition, R.S., Y.Y.; Supervision, Y.Y..

## Experimental Procedures

### Cell culture experiments

Two human iPS cell lines, 414C2tet-MyoD and 409B2tet-MyoD, established previously,^63^ were used in this study. Undifferentiated hiPSCs were seeded onto 6-well plates coated with iMatrix-511 silk (892021, Matrixome) and maintained with StemFit AK02N (Ajinomoto) at 37°C in a humidified atmosphere with 5% CO2. The differentiation of hiPSCs into myocytes was conducted as described.^64,65^ Briefly, hiPSC-414C2^tet-MyoD^ and hiPSC-409B2^tet-MyoD^ were plated onto 6-well plates pre-coated with Matrigel (1:100, 356231, Corning) in StemFit culture medium supplemented with 10 μM Y27632 (036-24023, Fuji Film-Wako) and grown overnight. On day 1, the medium was changed to Primate ES Cell Medium (RCHEMD001, Reprocell) containing 10 μM Y27632. On day 2, the medium was switched to Primate ES Cell Medium supplemented with 1 μg/mL Dox (D5897, LKT Laboratories) for inducing the expression of MyoD. On day 3, the medium was replaced with a differentiation medium consisting of MEMα medium (135-15175, Fuji Film-Wako) supplemented with 5% KnockOut Serum Replacement (10828028, Gibco), 1 μg/mL Dox, 50 U/mL penicillin, and 50 mg/mL streptomycin (09367-34, Nacalai Tesque), and cells were incubated for an additional 2 to 3 days to complete the myocyte differentiation. hiPSC-MCs were then incubated with Dulbecco’s modified Eagle medium (DMEM)/high glucose (043-30085, Fuji Film-Wako) containing 5% horse serum (HS) (16050122, Gibco) for 24 h before starting treatments.

HeLa (obtained from RIKEN CELL BANK) and HEK293T (American Type Culture Collection, ATCC) cells were maintained in DMEM/high glucose supplemented with 7.5% fetal bovine serum (FBS) and 1% penicillin/streptomycin. RC-13 cells (ATCC) were cultured in RPMI-1640 (189-02025, Wako) with 7.5% FBS and 1% penicillin/streptomycin. C2C12 myoblasts (ATCC) were maintained in DMEM/high glucose supplemented with 10% FBS and 1% penicillin/streptomycin. For differentiation into myotubes, C2C12 myoblasts were seeded into a 6-well plate 2 days before starting differentiation. On day 0, the medium was switched to DMEM with 2% HS, and cells were further incubated for 4–5 days to be differentiated into myotubes. The medium was changed every two days. All cells were maintained in a humidified atmosphere of 5% CO_2_ and 95% air at 37°C.

### Animal experiments

All animal experiments were approved by the Animal Care and Use Committee of the University of Tokyo, which is based on the Law for the Humane Treatment and Management of Animals (Law No. 105, 1 October 1973, as amended on 1 June 2020). Male C57BL6/J mice (4 or 7 weeks old) were purchased from Japan Clea (Tokyo, Japan) and acclimated for 1 week before experiments. All mice were housed in a temperature- and humidity-controlled room (23 ± 2°C, 55 ± 10% humidity) under a 12-hour dark/light cycle, with free access to a standard chow diet (Labo MR Stock, Nosan Corporation) and water.

To induce statin myopathy, 8-week-old male mice were randomly divided into 3 groups (n = 4–6 for each group). Mice were administered daily intraperitoneal injections with atorvastatin (30 mg/kg, A2476, Tokyo Chemical Industry), fluvastatin (30 mg/kg, F0820, Tokyo Chemical Industry), or saline (Control, D05352, Otsuka Pharmaceutical Factory) for 14 days. For investigating the role of the mevalonate pathway in vivo, 8-week-old male mice were randomly divided into four groups (n = 6 per group): Control (vehicle), Fluvastatin (30 mg/kg fluvastatin), Fluvastatin +MLN) (30 mg/kg fluvastatin and 100 mg/kg MLN), and Fluvastatin +GGOH (30 mg/kg fluvastatin and 50 mg/kg GGOH). Mice were treated daily as above for 14 days. On day 7 and 14, forelimb grip strength tests were conducted as described below.

For AAV-mediated knockdown experiments, 5-week-old male mice were used and randomly divided into different experiment groups (n = 5 per group). To silence FAM134B in the skeletal muscle, AAV vectors (either shScramble, shFAM134B#1, or shFAM134B#2 shRNAs, 5.0 × 10¹⁰ virus genomes, vg) were injected intramuscularly into both the left and right tibialis anterior (TA) muscles. Four weeks after virus injection, mice were subjected to daily treatment with either 30 mg/kg fluvastatin or saline for 14 consecutive days as above. Forelimb grip strength tests were conducted every week.

At the indicated time points, mice were anesthetized with isoflurane and subsequently euthanized by cervical dislocation. The quadriceps (Quad), gastrocnemius (Gas), tibialis anterior (TA), and soleus (Sol) from both legs were carefully removed, weighed, and photographed. The liver was also collected. These samples were then frozen in liquid nitrogen and stored at −80°C for subsequent experiments.

### RNA extraction and quantitative real-time PCR (qRT-PCR)

Total RNA from cells or animal tissues was extracted using ISOGEN (319-90211, Nippon Gene), and cDNA was synthesized using a High-Capacity cDNA Reverse Transcription Kit (43-688-14, Applied Biosystems) according to the manufacturer’s protocols. qRT-PCR was conducted using FastStart Universal SYBR Green Master (Roche Applied Science) on either a StepOnePlus Real-Time PCR System (Applied Biosystems) or a QuantStudio Flex Real-Time PCR System (Applied Biosystems). Target gene expression was normalized to the level of 18S ribosomal RNA, with expression in the control conditions set to 1. The sequence of qPCR primers used in this study is listed in Tables S1 and S2.

### RNA-sequencing and bioinformatic analysis

Total RNA of hiPSC-MCs from triplicated wells was pooled and submitted to Macrogen Japan. RNA sequencing was performed at Macrogen Japan Corp using NovaSeq 6000 system (Illumina).

The gene ontology (GO) information of human primary myotubes treated with statins was obtained from the NCBI Gene Expression Omnibus (GEO): GSE107998.^66^ Gene expressions in the muscle biopsies from statin tolerant and statin intolerant individuals was from GSE97254.^39^ Microarray data derived from multiple atrophic conditions were as follows: GSE48363 (gastrocnemius/plantaris muscles from mice with cancer cachexia),^67^ GSE220080 (quadriceps muscle from mice with spinal and bulbar muscular atrophy),^68^ GSE108299 (muscle from 48 h fasted mice), GSE80223 (soleus muscle of 30 days space-flown mice).^69^ Bioinformatic analysis and visualization were performed using R software (version 4.4.1). The edgeR bioconductor R package was used to analyze the differences in gene expression between the indicated samples and groups. Heatmap analysis was conducted the pheatmap R package. Venn diagram was analyzed and produced by VennDiagram R package. Gene-enriched pathways were annotated by reference to the Kyoto Encyclopedia of Genes and Genomes (KEGG) and visualized by ggplot2 R package.

### Plasmid construction

The FLAG-human SREBP1a (2-487) and FLAG-human SREBP2 (2-481) expression plasmids were previously described.^70^ FLAG-human FOXO1, FLAG-human FOXO3a, and FLAG-human FOXO4 expression plasmids were constructed using p3XFlag-CMV7.1 vector (Sigma). The coding sequence of mouse SERCA1 was cloned into p3XFlag-CMV-7.1 vector (Sigma) at the NotI/XbaI site to generate pFLAG-SERCA1. To generate EGFP-SERCA1 and pmCherry-EGFP-SERCA1 expression plasmids, coding region of mouse SERCA1 was first cloned into the pEGFP-C1 vector (Clonetech) at the XhoI/SacII sites, and then EGFP-SERCA1 was subcloned into pmCherry-C1 vector (Clonetech) at SacII/XmaI sites. The coding sequence of mouse FAM134B-S cDNA was cloned into p3XFlag-CMV-7.1 vector (Sigma) at the HindIII/EcoRI site to generate pFLAG-FAM134B-S, or cloned into pmCherry-C1 vector at the HindIII/EcoRI site to generate pmCherry-FAM134B-S. The coding sequence of human FAM134B-L was cloned into p3XFlag-CMV-14 vector (Sigma) at the HindIII/BamHI site to generate pFAM134B-L-FLAG. The human FAM134B-long promoter (-3000 to 0) and human FAM134B-short promoter (-3100 to -100) were cloned into pGL4.10[luc2] vector (Promega) at the KpnI/HindIII and KpnI/XhoI sites, respectively. 2xIRE sequence was cloned into pGL4.10[luc2] vector. The primers for constructing the above plasmids were listed in Table S3.

### Plasmid and siRNA transfection

Plasmid transfection with HeLa and C2C12 cells were performed using HilyMax (342-91103, Wako) according to the manufacturer’s instructions. Polyethylenimine Max (PEI-MAX, 24765, Polysciences) and ViaFect Transfection Reagent (E4981, Promega) were used for plasmid transfection in HEK293T and RC13 cells, respectively. Forty-eight hours post-transfection, cells were harvested for further experiments.

Scramble siRNA (siCtrl) and siRNA targeting FAM134B (siFAM) were purchased from Dharmacon (ON-TARGETplus Non-targeting Pool; ON-TARGETplus human RETREG1 siRNA – SMARTpool, 54463). hiPSCs were plated in a 6-well plate as above, and on day 1, siCtrl or siFAM (25 nM each) was transfected using Lipofectamine RNAiMAX Transfection Reagent (13778150, ThermoFisher Scientific) according to the manufacturer’s instructions. Differentiation into myocytes were conducted as described above. FAM134B siRNA nucleotide sequences were listed in Table S4.

### Luciferase reporter assay

HEK293T cells were seeded into a 12-well plate and cultured overnight. On day 1, reporter plasmid (200 ng/well), pCMV-β-galactosidase (100 ng/well), and expression plasmid of interest (500 ng/well) were transfected as above. Twenty-four hours post-transfection, a luciferase reporter assay was performed as described.^71^ Luciferase activity was normalized to β-galactosidase activity.

### Immunoblotting

Cell lysate was prepared using urea buffer (50 mM Tris–HCl pH 8.0, 50 mM sodium phosphate pH 8.0, 100 mM NaCl, 8 M urea) supplemented with phenylmethanesulfonyl fluoride (PMSF) (Sigma-Aldrich), 0.5% protease inhibitor cocktail (Nacalai Tesque), and 0.5% phosphatase inhibitor cocktail (Sigma-Aldrich).^72^ Cell lysates were scraped, vortexed for 30 min at 4°C, and spun at 20,630 xg for 10 min at 4°C, and the supernatants were collected as whole-cell lysates and stored at -80°C until further use. Tissue homogenates were prepared by homogenizing tissues in ice-cold radioimmunoprecipitation assay (RIPA) lysis buffer (50 mM Tris-HCl, pH 7.4, 150 mM NaCl, 1 mM EDTA, 1% Nonidet P-40, and 0.25% sodium deoxycholate) supplemented with PMSF, protease inhibitor cocktail, and phosphatase inhibitor cocktail as described.^65^ In brief, samples were left on ice for 10 min after being subjected to 20 strokes with a 25-gauge needle, followed by centrifugation at 20,630 xg for 10 min at 4°C. The supernatant was collected as tissue homogenate. Protein concentration of cell lysate and tissue homogenate was measured by BCA Protein Assay (A65453, Thermo Fisher) with BSA as a standard. Equal amounts of proteins were subjected to SDS-PAGE and immunoblot analysis according to a standard protocol. Primary antibodies used in this study were as follows; anti-FAM134B (21537-1-AP; 1:1,000) and anti-KTN1 (19841-1-AP; 1:1,000) from Proteintech; anti-GAPDH (2118; 1:1,000), anti-LC3B (2775;1:1,000), anti-SERCA1 (4219; 1:1,000), anti-Calnexin (2679; 1:1,000), and anti-GFP (2956; 1:1,000) from Cell Signaling Technology; anti-Atrogin-1 (ab168372; 1:1,000) from Abcam; anti-FLAG, Clone M2 (F1804; 1:1,000), anti-β-Actin (A5441; 1:1,000) from Sigma-Aldrich; anti-RyR (sc-376507; 1:1,000), anti-SERCA2 (sc-376235; 1:1,000), and anti-HSP90 (sc-13119; 1:1,000) from Santa Cruz Biotechnology; anti-RFP (PM005; 1:1,000) from MBL. Second antibodies were HRP-linked anti-mouse IgG (7076; 1:5,000), and HRP-linked anti-rabbit IgG (7074; 1:5,000) from Cell Signaling Technology. The signal was detected by a FUSION SOLO S chemiluminescence imaging system (Vilber-Lourmat) using the Amersham ECL Immunoblotting detection reagents (GE Healthcare). Band intensities were quantified using ImageJ software or Evolution-Capt software (Vilber Lourmat). GAPDH, β-Actin or HSP90 was used as an internal control for normalization.

### Immunoprecipitation (IP)

IP was performed according to a protocol previously described.^73^ Cells were lysed by passing through a 25-gauge needle 20 times in IP lysis buffer (25 mM Tris-HCl pH 7.4, 150 mM NaCl, 1 mM EDTA, 5% glycerol, and 0.5% Nonidet P-40, PMSF, protease inhibitor cocktail, and phosphatase inhibitor cocktail). After centrifugation at 20,630 xg for 10 min, the supernatant was incubated with pre-cleared rabbit IgG agarose beads (A2909, Sigma-Aldrich) for 1 h with rotation at 4 °C. Afterward, the supernatant was incubated with a primary antibody against FAM134B (83414; 1:200, Cell Signaling Technology) or control rabbit IgG antibody (PM035; 1:200, MBL) overnight with rotation at 4 °C, and then incubated with pre-cleared protein A-Agarose (11719408001, Roche) for 1 h with rotation at 4 °C. After washing with IP lysis buffer, IP products were eluted with sample buffer for immunoblotting analysis. For IP of FLAG-tagged proteins, cell lysate was pre-cleared by incubating with mouse IgG-agarose (A0919, Sigma-Aldrich) for 1 h at 4 °C. IP was then performed using anti-FLAG M2 affinity gel (A2220, Sigma-Aldrich) according to the manufacturer’s instructions. The precipitated proteins were eluted using 3× FLAG peptides (F4799, Sigma-Aldrich) and boiled in sample buffer for immunoblotting.

### Immunofluorescence staining and image analysis

For immunofluorescence staining, cells were seeded onto an 18 mm x 18 mm coverslip (Matsunami) except hiPSCs, where these cells were seeded onto a 35 mm-film-bottom dish (FD10300, Matsunami). For labeling autophagosomes and autolysosomes, cells were incubated with 0.3 μM or 0.1 μM DAPGreen (340-09291, DOJINDO) in culture medium for 30 min at 37°C as described.^37^ To monitor the lysosomes, cells were incubated with 50 nM LysoTracker Deep Red (L12492, Thermo Fisher) at 37 °C for 2 h. Cells were fixed with 4% paraformaldehyde (PFA, 163-20145, Wako) in PBS, washed with PBS, and then permeabilized with 0.1% Triton X-100 in PBS for 5 min at room temperature. Cells were blocked with 5% FBS (in PBS) for 1 h and incubated with primary antibodies at a specific concentration (see below) for 1 h at room temperature. After washing with PBS, specimens were incubated with Alexa Fluor 488–conjugated anti-mouse IgG, Alexa Fluor 488–conjugated anti-rabbit IgG, Alexa Fluor 568–conjugated anti-mouse IgG, Alexa Fluor 568–conjugated anti-rabbit IgG or Alexa Fluor 647–conjugated anti-mouse IgG secondary antibodies (Thermo Fisher, 1:500) for 1 h at room temperature, washed with PBS, and mounted with ProLong Diamond Antifade Mountant with DAPI (P36971, ThermoFisher). Primary antibodies used were as follows: anti-KDEL (M181-3; 1:400) from MBL, anti-myosin heavy chain (MAB4470; 1:100) from R&D Systems; anti-FAM134B (83414; 1:200), and anti-SERCA1 (4219; 1:200) from Cell Signaling Technology; anti-FLAG M2 (F1804; 1:400) from Sigma-Aldrich.

For immunofluorescent staining of mouse TA muscles, tissue cryosections were fixed with 4% paraformaldehyde for 15 minutes, permeabilized for 10 minutes with 0.3% Triton X-100 in PBS, and then blocked for 1 h at room temperature in a blocking buffer (0.1% Triton X-100 and 5% FBS in PBS). Primary antibodies were incubated overnight at 4°C as follows; anti-KDEL (M181-3; 1:200, MBL), anti-FAM134B (83414; 1:200, Cell Signaling Technology), and anti-Dystrophin (sc-73592; 1:200, Santa Cruz Biotechnology). The appropriate secondary antibodies conjugated with Alexa Fluor 488 or Alexa Fluor 568 were diluted in 2% blocking buffer, and specimens were incubated for 1 h at room temperature. Sections were mounted with ProLong Diamond Antifade Mountant with DAPI.

Confocal images were acquired under a Zeiss LSM 800 confocal laser scanning microscope (Carl Zeiss) with a Plan-Apochromat 63x/1.40 Oil DIC M27 objective or a Plan-Apochromat 40x/1.40 Oil DIC M27 objective (Carl Zeiss). Images were processed with Zen software (Zeiss). For measuring myocyte diameters, images were obtained using a fluorescence microscope BZ-X810 (KEYENCE) with a CFI Plan Fluor 10x/0.30 objective (Nikon). Myocyte diameters were quantified as previously described^64,74^ by using IC Measure (The Imaging Source). Briefly, the diameters of at least 50 MHC-positive myocytes were measured at three points (the thickest point and two points 50 µm apart from the thickest point) per myocyte, and the average diameter of each myocyte was calculated.

### Transmission electron microscopy

hiPSC-MCs were cultured on 18 mm x 18 mm coverslip (Matsunami) in a 6-well plate. Cells were fixed with 2.5% glutaraldehyde in 0.1 M phosphate buffer at 4 °C overnight, and washed with 0.1 M phosphate buffer, and postfixed with 1% OsO4 in 0.1 M phosphate buffer at 4 °C for 1 h. After dehydration in a graded ethanol series (70%, 90%, 95%, 99.5%, and 100%), ultrathin cell sections were stained with uranyl acetate and the Lead stain solution (18-0875, Sigma-Aldrich) and observed using a transmission electron microscope (JEM-1400Plus, JEOL) at an acceleration voltage of 100 kV with a Keen view CCD camera (Olympus Soft Imaging Solution).

### AAV production

The recombinant AAV2/8-GFP-U6 vector was reassembled and produced from the previously established recombinant AAV2/8-thyroxine binding globulin (TBG)-GFP vector and the pAAV2/1-GFP-U6 plasmid (gifted from Dr. Eguchi, The Institute of Medical Science, The University of Tokyo). In brief, the GFP-U6 promoter region was amplified from the pAAV2/1-GFP-U6 plasmid using PacI/HindIII cloning sites and replaced the TBG-GFP promoter region in rAAV2/8-TBG-GFP at the same cloning sites as described.^75^ The shRNA sequences targeting FAM134B, including shFAM134B#1 and shFAM134B#2, were inserted between the BamHI and EcoRI cloning sites driven by the U6 promoter. The rAAV-8 viral vector for triggering FAM134B knockdown was packaged and harvested from HEK293T cells using a triple-plasmid co-transfection method, followed by purification through cesium chloride (CsCl) density gradient ultracentrifugation as reported.^76^ The rAAV2/8-TBG-GFP vector plasmid, and packaging plasmids, including p5E18-VD2/8 capsid and pXX6-80 helper, were generously provided by Dr. A. Miyajima and Dr. T. Itoh (Institute of Molecular and Cellular Biosciences, University of Tokyo). AAV titers were determined by Taqman-based qRT-PCR as reported previously.^77^ The PCR primers used for shRNA sequences and AAV titration are listed in Tables S4 and S5.

### Grip strength test

The grip strength of the mice was measured using a grip strength meter (MK-380V, Muromachi) as described previously.^75^ A mouse was placed on the metal mesh of the meter with all four paws, and its tail was gently pulled back horizontally until the grip was broken and the peak force measurement was recorded. Grip strength tests were repeated six times with a 30-sec interval for each mouse. The grip strength was calculated as the average of six trials and normalized to body weight.

### Histology of muscle cryosections

Muscle tissues were carefully dissected and frozen in pre-cooled liquid isopentane in liquid nitrogen, embedded in Tissue-Tek O.C.T. compound (Sakura Finetek) and stored at −80 °C until processed. Frozen muscle samples were cut into 10 µm cryosections using a cryostat (NX70, Thermo Fisher) at −20 °C then mounted on adhesive glass slides (0863141690, MATSUNAMI). Muscle cryosections were stained with Hematoxylin and Eosin (H&E) according to a standard protocol and observed with a fluorescence microscopy (BZ X-800, Keyence). Quantification of myofiber cross-section area (CSA) was performed by Cellpose segmentation combined with Fiji plugin LabelsToRois as previously described.^78^

### Statistical analysis

Results are presented as mean ± SD from at least three or more independent biological replicates. All experiments were repeated on at least three occasions with similar results, except in vivo studies where at least two similar experiments were performed. Statistical analyses were performed by unpaired Student’s t-test or one-way ANOVA with Turkey-Kramer or Dunnett post-hoc tests using Prism 10 software (GraphPad), as specified in figure legends. p values less than 0.05 were considered statistically significant.

## Abbreviations

HMG-CoA: 3-hydroxy-3-methylgrutaryl coenzyme A
AAV: Adeno-associated virus
CVD: Cardiovascular disease
ER: Endoplasmic reticulum
GGOH: Geranylgeraniol
GGPP: Geranylgeranyl pyrophosphate
HMGCR: Gastrocnemius
Gas: HMG-CoA reductase
hiPSCs: Human induced pluripotent stem cells
hiPSC-MCs: hiPSC-derived myocytes
LDL: Low-density lipoprotein
MVA: Mevalonate
MLN: Mevalonolactone
Quad: Quadriceps
SAMS: Statin-associated muscle symptoms
SIM: Stain-induced myopathy
SREBP: Sterol regulatory element-binding protein
SR: Sarcoplasmic reticulum
SERCA1: Sarcoplasmic reticulum calcium ATPase 1
TA: Tibialis anterior
Sol: Soleus

