## Supplemental data for "Selective induction by statins of FAM134B-mediated sarcoplasmic reticulum (SR)-phagy degrades the SR calcium pump SERCA1 and contributes to myopathy"


Supplemental Tables: Table S1–S5
Supplemental Figures: Figure S1–S8

**Table S1. Primers used for qRT-PCR analysis of human genes.**

| Gene | Primer sequences |
|---|---|
| *FAM134B-L* | Fw: 5’-AACCTGCTGTTCTGGTTCCTT-3’<br>Rv: 5’-TCACTGAGGCTTCTCCACAAC-3’ |
| *FAM134B-S* | Fw: 5’-TGCAGCCTTTGCCACTGTTA-3’<br>Rv: 5’-GCCTGGGTCTTTCATCTGGT-3’ |
| *FAM134A* | Fw: 5’-CTGACGTTTCAGCATCATCC-3’<br>Rv: 5’-GAACATAGTGTCCCAACACAGC-3’ |
| *FAM134C* | Fw: 5’-GCTGGACGATTCTACTGTTGCC-3’<br>Rv: 5’-GCCTTCCGTTAGAGGTGTTTGG-3’ |
| *CCPG1* | Fw: 5’-TATCCAGCTTTGGAGGAAACC-3’<br>Rv: 5’-AGAGGAAGAGCCCATGTTAAAG-3’ |
| *TEX264* | Fw: 5’-CCAGGAAGACCAGATCCATTTC-3’<br>Rv: 5’-GGGCTCACTTCCAAGCTTAC-3’ |
| *SEC62* | Fw: 5’-GGATCTATGACCCAGTTCACTTT-3’<br>Rv: 5’-GGTGCCTTCCTCCAGTTATG-3’ |
| *RTN3L* | Fw: 5’-TAGATGCTGTTTCCAGCCTTAG-3’<br>Rv: 5’-AGAGCCAGGATGAGGTAAGA-3’ |
| *ATL3* | Fw: 5’-GAAGATGGGTGGGAAGGATTT-3’<br>Rv: 5’-GGCTACAACCTCAAGACCTATG-3’ |
| *CALCOCO1* | Fw: 5’-GAGGCTGAAGATGAGAAGTCAG-3’<br>Rv: 5’-GGAAAGCGCTCCTTACAGATAG-3’ |
| *HMCGS1* | Fw: 5’-GACTTGTGCATTCAAACATAGCAA-3’<br>Rv: 5’-GCTGTAGCAGGGAGTCTTGGTACT-3’ |
| *SERCA1* | Fw: 5’-GGTGCTGGCTGACGACAACT-3’<br>Rv: 5’-AAGAGCCAGCCACTGATGAG-3’ |
| *SERCA2* | Fw: 5’-GTGCTCCTGAAGGTGTCATT-3’<br>Rv: 5’-GAGTCCTCAAGGTGCATTTCT-3’ |
| *RYR* | Fw: 5’-ATCCTGACTGAAGACCACAGTT-3’<br>Rv: 5’-AGCATAGGCCATGTACAGGTAA-3’ |
| *DNM2* | Fw: 5’-GAGTTTGACGAGAAGGACTTA-3’<br>Rv: 5’-GATTAGCTCCTGGATAACCAG-3’ |
| *MTM1* | Fw: 5’-TGGAAGAATACAGGAGGC-3’<br>Rv: 5’-TGGAATTCGATTTCGGGAC-3’ |

**Table S2. Primers used for qRT-PCR analysis of mouse genes.**

| Gene | Primer sequences |
|---|---|
| *Fam134b-L* | Fw: 5’-CTGCTCACCTTCCTGGGTG-3’ |
| | Rv: 5’-TTCCCAGCTTTCAGTGAGGC-3’ |
| *Fam134b-S* | Fw: 5’-TGTGGACCAGGCAGAAG-3’ |
| | Rv: 5’-GCAGACCAGGAGGCAAA-3’ |
| *Hmgcs1* | Fw: 5’-GCGTCTTTGCTTGTGTGTAATG-3’ |
| | Rv: 5’-GCGTCTTTGCTTGTGTGTAATG-3’ |
| *Ldlr* | Fw: 5’-TGTGAAAATGACTCAGACGAACAA-3’ |
| | Rv: 5’-GGAGATGCACTTGCCATCCT-3’ |
| *Fbxo32/Atrogin-1* | Fw: 5’-CCTGCCTGTGTGCTTACAACTG-3’ |
| | Rv: 5’-GGTCCCGCCCGTCACT-3’ |
| *Trim63/MuRF1* | Fw: 5’-ACCTGCTGGTGGAAAACATC-3’ |
| | Rv: 5’-CTTCGTGTTCCTTGCACATC-3’ |
| *Map1lc3b* | Fw: 5’-CCCCACCAAGATCCCAGT-3’ |
| | Rv: 5’-CGTTCATGTTCACGTGGT-3’ |
| *Ccpg1* | Fw: 5’-CCCCACCAAGATCCCAGT-3’ |
| | Rv: 5’-CGTTCATGTTCACGTGGT-3’ |
| *Atl3* | Fw: 5’-AAGATCTGCCTCACCCCAAGTC-3’ |
| | Rv: 5’-CTCCCCCACAAACCTCTTCCAT-3’ |
| *Tex264* | Fw: 5’-TCCAGTAGCGATGACACCAG-3’ |
| | Rv: 5’-CTGGATTGTGCCATAGAAATGTC-3’ |
| *Serca1* | Fw: 5’-GGTACTGGCCGATGATAACT-3’ |
| | Rv: 5’-AAGAGCCAGCCACTGATAAG-3’ |
| *Serca2* | Fw: 5’-TGACGGCGGTCCAAGAGTCT-3’ |
| | Rv: 5’-CATGGACAAGCAGATGGAGC-3’ |
| *Ryr1* | Fw: 5’-TACAAAGCCTCCTGGATCCT-3’ |
| | Rv: 5’-AGCGTAAGCCATGTATAGGT-3’ |
| *Dnm2* | Fw: 5’-GCAGGAGATCGAAGCAGAGAC-3’ |
| | Rv: 5’-GGTCGATGAGGGTCAAGTTCAA-3’ |
| *Mtm1* | Fw: 5’-CATGCGTCACTTGGAACTGTGG-3’ |
| | Rv: 5’-GCAATTCCTCGAGCCTCTTT-3’ |
| *18S rRNA* | Fw: 5’-ACCGCAGCTAGGAATAATGGA-3’ |
| | Rv: 5’-GCCTCAGTTCCGAAAACCA-3’ |

**Table S3. Sequences of the primers used for cloning.**

| Gene | Primer sequences |
|---|---|
| Human | |
| FAM134B-long promoter | Fw: 5’-GTGAGTGGCCTCTGTTAGAGAG-3’<br>Rv: 5’-CTTCAGCTGTGCTTCCAGACAG-3’ |
| FAM134B-short promoter | Fw: 5’-CCTCCACCATGATTCTAAGTTTCC-3'<br>Rv: 5’-ACCCAGTAGAGACGTTCTTTCC-3’ |
| pFLAG-SREBP1a-mature | Fw: 5’-ATAAGAATGCGGCCGCATGGACGAGCCACCCTTCAGC-3’<br>Rv: 5’-GCTCTAGACTAGCAGGGGCAGTGGCAGCGGT-3’ |
| pFLAG-SREBP2-mature | Fw: 5’-ATAAGAATGCGGCCGCGACGACAGCGGCGAGCTG-3'<br>Rv: 5’-GCTCTAGACGGATTCTTCTGTGTGTCCTC-3’ |
| pFAM134B-long-FLAG | Fw: 5’-AATTAAGCTTATGGCGAGCCCGGCGCCTC-3'<br>Rv: 5’- AATTGGATCCATGGCCTCCCAGCAGATTT-3’ |
| pFLAG-FAM134B-short | Fw: 5’-ATAAAGCTTGCCACCATGCCTGAAGGTGAAGACTTTGGAC-3'<br>Rv: 5’- ATATGAATTCTTAATGGCCTCCCAGCAGATTTGA-3’ |
| pmCherry-FAM134B-short | Fw: 5’- ATAAAGCTTATGCCTGAAGGTGAAGACTTTGGAC-3'<br>Rv: 5’-ATATGAATTCTTAATGGCCTCCCAGCAGATTTGA-3’ |
| Mouse | |
| pFLAG-SERCA1 | Fw: 5’-AGCGGCCGCAGCCACCATGGAGGCCGCGCACTCCAAGT-3’<br>Rv: 5’-ATTCTAGATTATCCCTCCAGATAGTTCCGAGC-3’ |
| pEGFP-SERCA1 | Fw: 5’-ATATCTCGAGAAGCCACCATGGAGGCCGCGCACTCCAAGT-3’<br>Rv: 5’-ATATCCGCGGTTATCCCTCCAGATAGTTCCGAGC-3’ |
| pmCherry-eGFP-SERCA1 | Fw: 5’-ATATCCGCGGAAGTGAGCAAGGGCGAGGAGCTG-3’<br>Rv: 5’-ATATCCCGGGTTATCCCTCCAGATAGTTCCGAGC-3’ |
| pFAM134B-long-FLAG | Fw: 5’-AATTAAGCTTATGGCGAGCCCGGCGCCTC-3'<br>Rv: 5’- AATTGGATCCATGGCCTCCCAGCAGATTT-3’ |
| pFLAG-FAM134B-short | Fw: 5’-ATAAAGCTTGCCACCATGCCTGCAGGCGGCGGCTGT-3'<br>Rv: 5’- ATATGAATTCTTAATGGCCTCCAAGCAGATTGGA-3’ |

**Table S4. Sequences of the primers for target siRNA and shRNAs.**

| Oligonucleotides | Sequences |
|---|---|
| siFAM134B | Sequence 1: 5’-AGUUGUAGACUUAGGCUUA-3' |
| | Sequence 2: 5’-UCAGAAGAAACGUGAGAGA-3’ |
| | Sequence 3: 5’-GAGAGUGAAUUGGGACUUA-3’ |
| | Sequence 4: 5’-CCGAAAUUAUCAAGGUAUC-3' |
| shFAM134B#1 | Fw: 5’-GATCCGAGGTATCCTGGACTGATAATCTCGAGATTATCAGTCCAGGATACCTCTTTTTG-3’ |
| | Rv: 5’-AATTCAAAAAGAGGTATCCTGGACTGATAATCTCGAGATTATCAGTCCAGGATACCTCG-3’ |
| shFAM134B#2 | Fw: 5’-GATCCCACAAGGATGACAGTGAATTACTCGAGTAATTCACTGTCATCCTTGTGTTTTTG-3’ |
| | Rv: 5’-AATTCAAAAACACAAGGATGACAGTGAATTACTCGAGTAATTCACTGTCATCCTTGTGG-3’ |

**Table S5. Sequences of the probe primers for AAV titration.**

| Primer | Sequences |
|---|---|
| Probe | 5’-FAM-TGCTGACGCAACCCACTGGT-BHQ-3' |
| *Forward* | 5’-CCGTTGTCAGGGCAACGTG-3’ |
| *Reverse* | 5’-AGCTGACAGGTGGTGGCAAT-3’ |

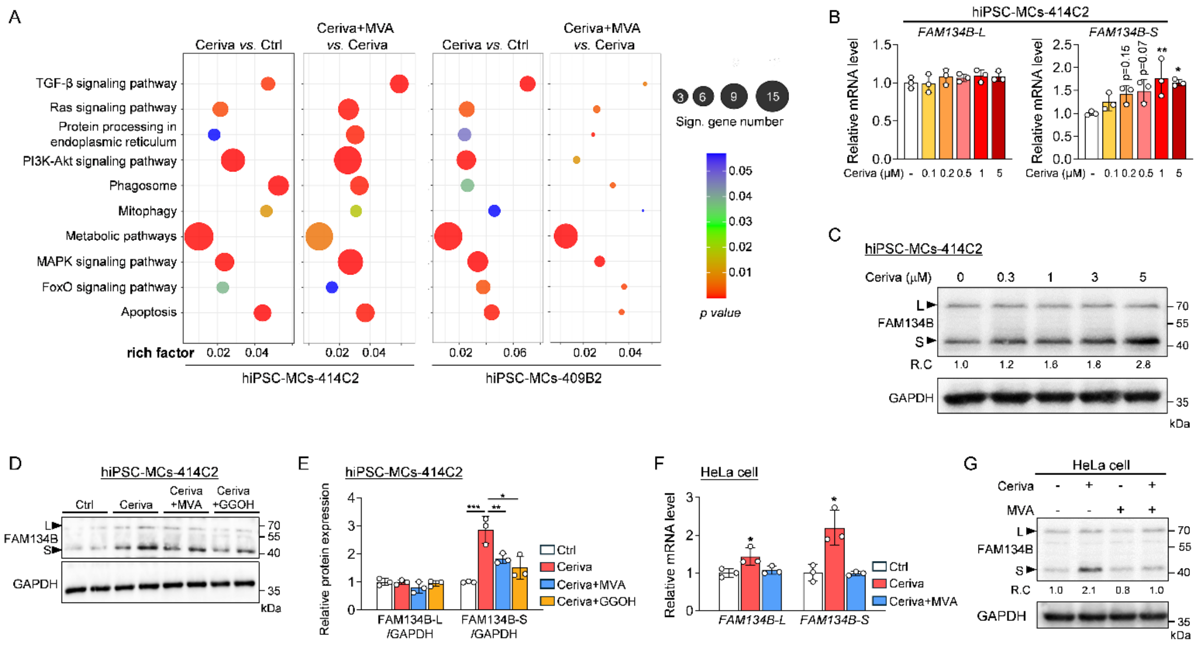

**Figure S1. Cerivastatin selectively upregulates FAM134B-S expression in a mevalonate pathway-dependent manner.**

(A) RNA-seq analysis was conducted in hiPSC-MCs-414C2 and hiPSC-MCs-409B2 treated with or without 5 μM cerivastatin and 200 μM MVA for 16 h. Bubble chart showing enriched KEGG pathways of differentially expressed genes in hiPSC-MCs. The color and size of the dots represent the range of the p-value and the number of genes mapped to each KEGG pathway, respectively.

(B) hiPSC-MCs (414C2) were treated with the indicated concentration of cerivastatin for 16 h. mRNA expression of *FAM134B-L* and *FAM134B-S* were analyzed. n=3.

(C) FAM134B expression was assessed by immunoblotting in hiPSC-MCs-414C2 treated as above. GAPDH was used as an internal control. Relative changes (RCs) of FAM134B-S expression are shown at the bottom of the FAM134B blot.

(D, E) Immunoblot images (D) and quantification data (E) of FAM134B in hiPSC-MCs-414C2. GAPDH was used as an internal control. n=3 per group for quantification.

(F) HeLa cells were treated with 5 μM cerivastatin in the presence or absence of 200 μM MVA for 16 h. mRNA expression of *FAM134B-L* and *FAM134B-S* were analyzed. n=3.

(G) FAM134B expression was assessed by immunoblotting in HeLa cells treated as above. GAPDH was used as an internal control. Relative changes (RCs) of FAM134B-S expression are shown at the bottom of the FAM134B blot.

Data shown as mean ± S.D. Statistical analyses were performed by one-way ANOVA with a Dunnett post hoc test. *, $p < 0.05$, **, $p < 0.01$, ***, $p < 0.001$ vs Control group (B), or vs cerivastatin-treated group (E and F).

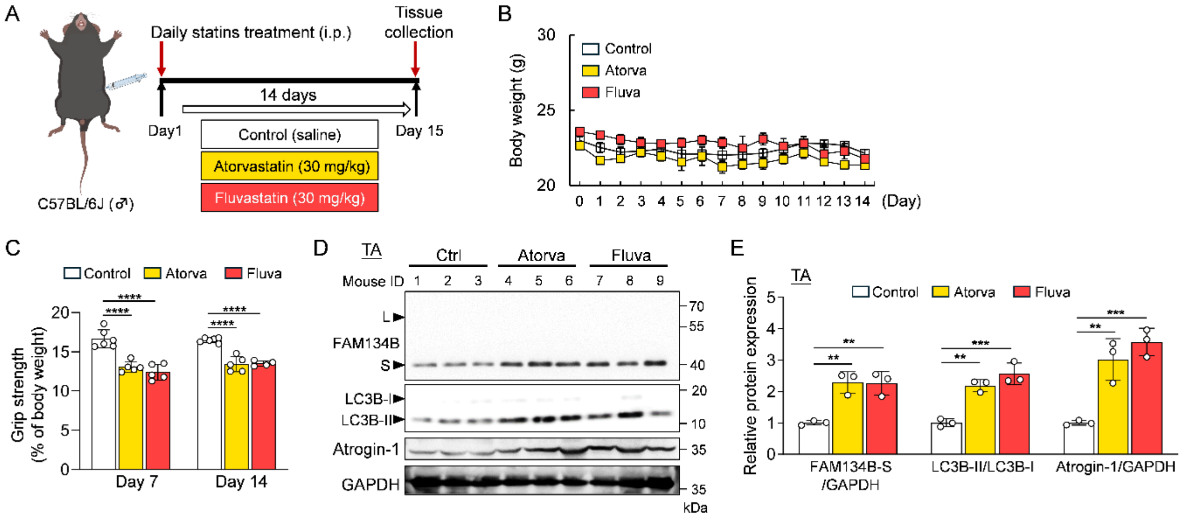


**Figure S2. Establishment of statin-induced myopathy model in mice.**

(A) Schematic diagram of experimental design. Mice were treated with or without atorvastatin and fluvastatin for 14 days.

(B) Body weight was measured daily during the treatments (n=4–6).

(C) Grip strength was measured on days 7 and 14 and normalized to body weight (n=4–6).

(D, E) Protein expression of FAM134B, LC3B, and Atrogin-1 in TA (D). The intensity of bands normalized to GAPDH was quantified (E) (n=3).

Data shown as mean ± S.D. (n = 4–6). Statistical analyses were performed by one-way ANOVA with Dunnett post hoc test. **, $p < 0.01$, ***, $p < 0.001$, ****, $p < 0.0001$ vs Control.

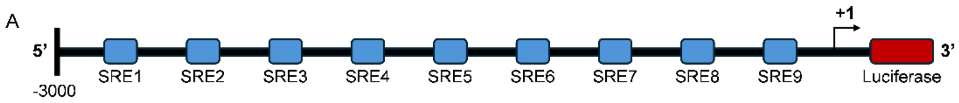


| Name | Position | Sequence |
|---|---|---|
| SRE1 | -2744/-2735 | gtcatgccag |
| SRE2 | -2433/-2420 | tgaacacgtgaact |
| SRE3 | -1906/-1897 | ctcactccag |
| SRE4 | -1448/-1439 | ctcagcctat |
| SRE5 | -1232/-1223 | ctcgggtgat |
| SRE6 | -1099/-1086 | ggctcaagtgattc |
| SRE7 | -756/-747 | atcagttcat |
| SRE8 | -434/-425 | atcagcctat |
| SRE9 | -305/-296 | gtggggttat |

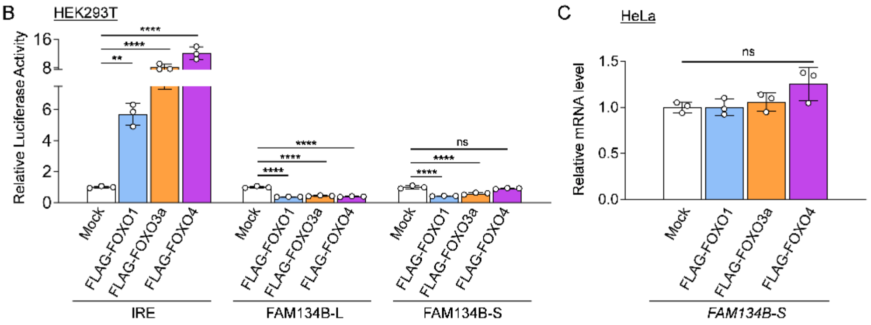


**Figure S3. FOXOs indirectly regulate FAM134B-short isoform.**

(A) Schematic representation of partial nucleotide sequence of human *FAM134B-S* gene proximal promoter [GenBank: NM_019000.5]. +1 represents transcriptional start site. Blue boxes indicate the putative binding sites for SREBP transcription factors. The position and sequence of putative SREs appear in the table below.

(B) Luciferase reporter assay was conducted in HEK293T cells. Cells were co-transfected with 2xIRE (as a positive control), human *FAM134B-L* or human *FAM134B-S* promoter gene luciferase reporter plasmid, along with either one of the three human FOXO expression plasmids (pFLAG-FOXO1, pFLAG-FOXO3a, and pFLAG-FOXO4). The luciferase activity was measured 24 h after transfection (n=3).

(C) FLAG-FOXO1, FLAG-FOXO3a, and FLAG-FOXO4 were forcedly expressed in HeLa cells. mRNA level of *FAM134B-S* was analyzed by RT-qPCR (n=3).

Data shown as mean ± S.D. Statistical analyses were performed by one-way ANOVA with a Dunnett post hoc test. **, $p < 0.01$, ****, $p < 0.0001$ vs mock.

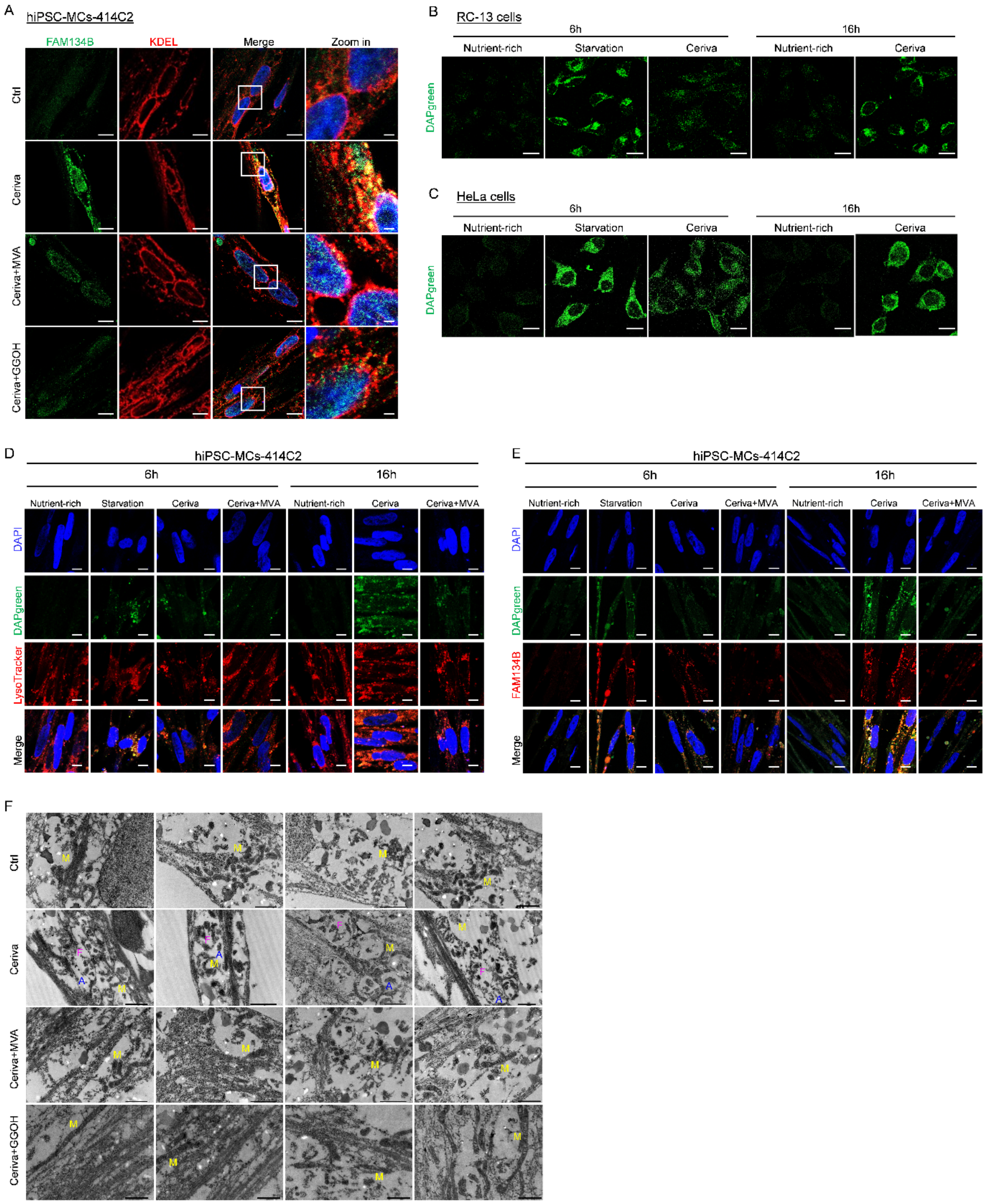


**Figure S4. Statin induces FAM134B-mediated ER-phagy.**

(A) Expression and localization of FAM134B in hiPSC-MCs-414C2 treated with 5 μM cerivastatin in the presence or absence of MVA (200 μM) or GGOH (100 μM) for 16 h. Shown are representative confocal images of cellular localization of FAM134B (green) and KDEL-positive ER (red) in hiPSC-MCs. Scale bar, 5 μm.

(B, C) RC-13 (B) and HeLa (C) cells were preincubated with DAPGreen (0.1 μM) for 30 min.

Cells were then starved in EBSS medium for 6 h, in the presence or absence of 1 μM cerivastatin for 6 h or 16 h, and analyzed using confocal microscopy. Representative images are shown. Scale bar, 5 μm.

(D) hiPSC-MCs-414C2 were preincubated with DAPGreen (0.1 μM) for 30 min. Then, cells were starved in EBSS medium for 6 h (as a positive control) or treated with cerivastatin (5 μM) in the presence or absence of 200 μM MVA for 6 or 16 h. LysoTracker (0.5 μM) were added 2 h before fixation. Representative confocal images of the localization of DAPGreen (green) and LysoTracker (red) are shown. Scale bar, 5 μm.

(E) Localization of DAPGreen (green) and FAM134B (red) in hiPSC-MCs-414C2 treated as above. Scale bar, 5 μm.

(F) TEM images of hiPSC-MCs-414C2 treated with 5 μM cerivastatin in the presence or absence of MVA (200 μM) or GGOH (100 μM) for 16 h. Scale bar, 1 μm. M, mitochondria; A, autolysosome; F, ER-like fragments.

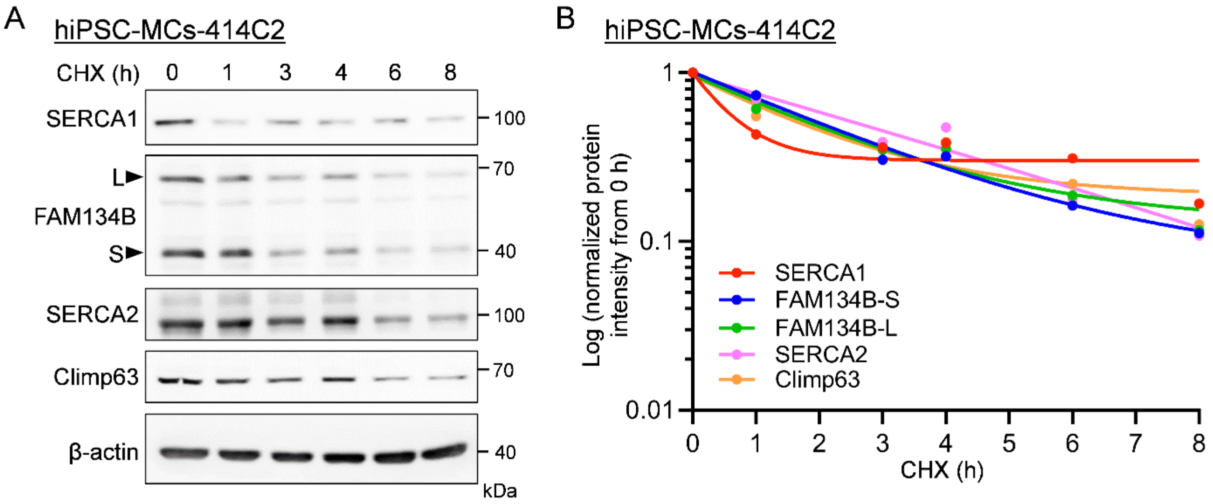


**Figure S5. Degradation rate of the ER/SR proteins in hiPSC-MCs.**

(A-B) hiPSC-MCs-414C2 were not treated or treated with CHX (50 μM) for 1, 3, 4, 6, 8 h as indicated. SERCA1, FAM134B, SERCA2 and Climp63 protein levels were assessed by immunoblotting with β-actin as an internal control (A). Relative changes of each protein were plotted (B).

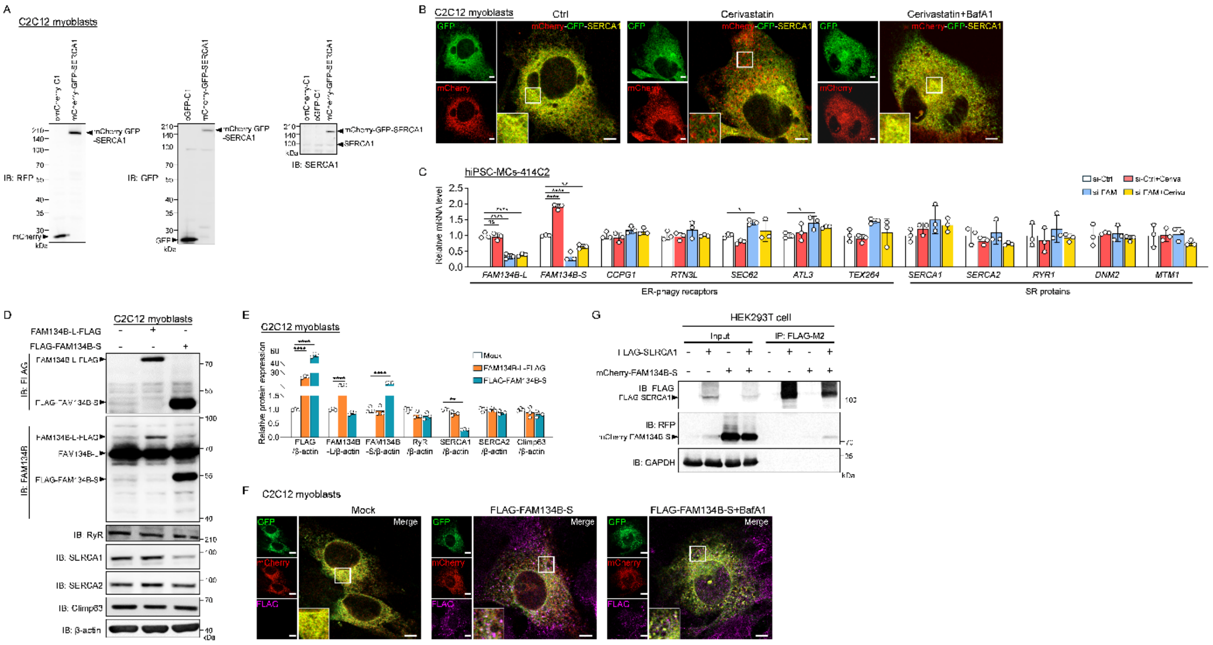

**Figure S6. SERCA1 protein is degraded by FAM134B-mediated SR-phagy.**

(A) mCheery-C1, EGFP-C1 and mCherry-eGFP-SERCA1 were forcedly expressed in C2C12 myoblasts. Protein expression of mCherry, GFP and SERCA1 was analyzed by immunoblot.

(B) mCherry-eGFP-SERCA1 was forcedly expressed in C2C12 myoblasts, and cells were treated with 5 μM cerivastatin for 16 h. BafA1 (100 nM) was added 3 h before fixation. Shown are representative confocal images of the localization of GFP (green) and mCherry (red). Scale bar, 5 μm.

(C) hiPSC-MCs-414C2 were transfected with siControl or siFAM134B (25 nM each). Cells were treated with or without 5 μM cerivastatin for 16 h. mRNA expression of ER-phagy receptors and SR/ER proteins was analyzed by RT-qPCR. n=3 per group.

(D, E) Protein expressions of FLAG, FAM134B, RyR, SERCA1, SERCA2, and Climp63 in C2C12 myoblasts overexpressed either FAM134B-L-FLAG or FLAG-FAM134B-S (D). β-actin was used as an internal control, and the expression of indicated proteins was quantified (E) (n=3).

(F) mCherry-eGFP-SERCA1 and FLAG-FAM134B-S were forcedly expressed in C2C12 myoblasts, and cells were treated with 100 nM BafA1 for 3 h before fixation. Shown are representative confocal images of the localization of GFP (green), mCherry (red) and FLAG-FAM134B-S (magenta). Scale bar, 5 μm.

(G) FLAG-SERCA1 and mCherry-FAM134B-S were co-expressed in HEK293T cells for 48h. Immunoprecipitation was performed with anti-FLAG M2 as described in Methods. FLAG-SERCA1 and mCherry-FAM134B-S were detected by immunoblot with anti-FLAG and anti-

RFP antibodies, respectively. GAPDH was used as an internal control.

Data shown as mean ± S.D. Statistical analyses were performed by one-way ANOVA with a Tukey-Kramer (C) or Dunnett post hoc test (E). *, $p < 0.05$, **, $p < 0.01$, ****, $p < 0.0001$.

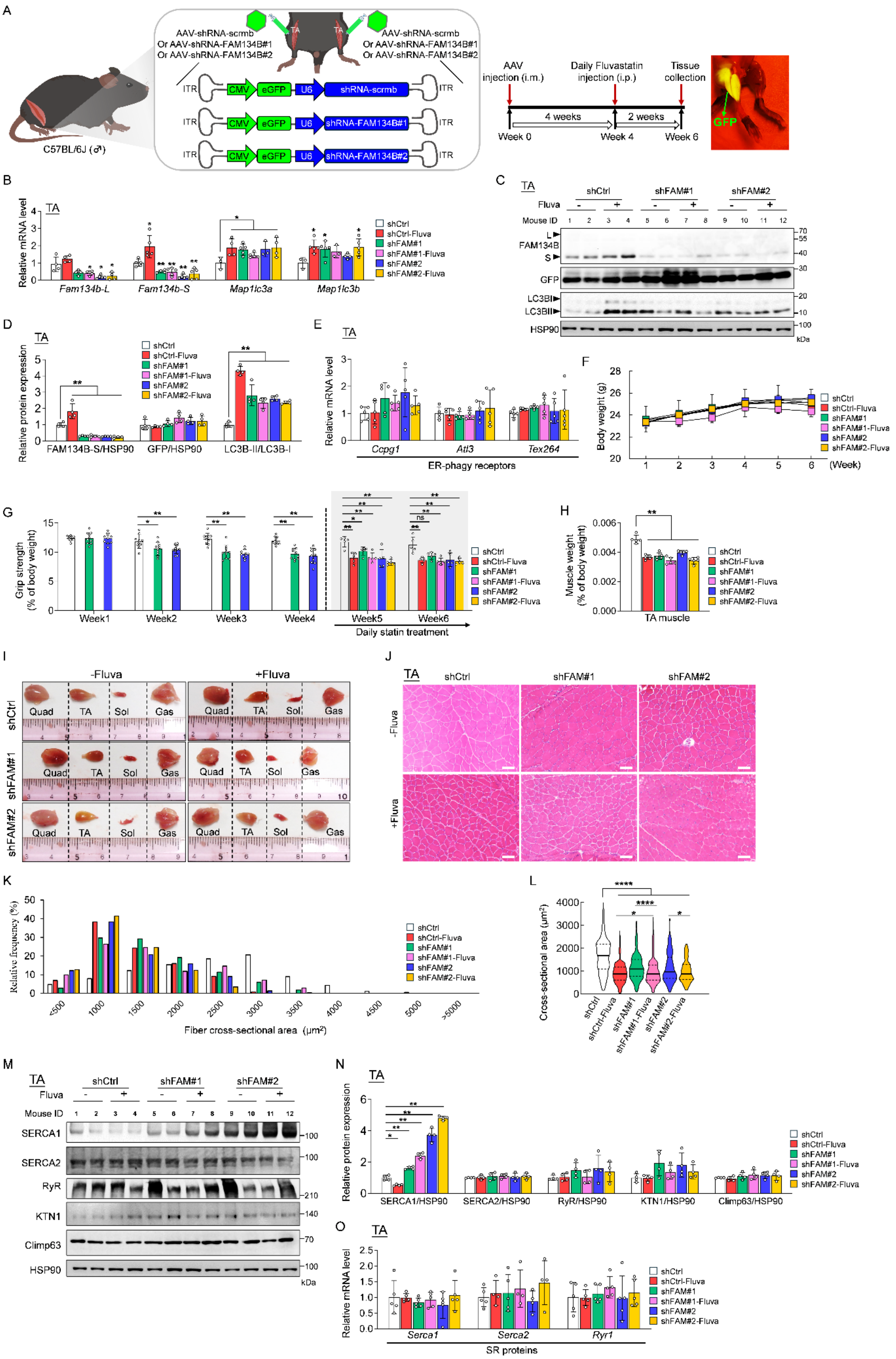
A
AAV-shRNA-scrmb
Or AAV-shRNA-FAM134B#1
Or AAV-shRNA-FAM134B#2
C57BL/6J (♂)
ITR CMV eGFP U6 shRNA-scrmb ITR
ITR CMV eGFP U6 shRNA-FAM134B#1 ITR
ITR CMV eGFP U6 shRNA-FAM134B#2 ITR
AAV injection (i.m.)
Daily Fluvastatin injection (i.p.)
Tissue collection
4 weeks
2 weeks
Week 0
Week 4
Week 6
GFP
B
TA
Relative mRNA level
Fam134b-L
Fam134b-S
Map1lc3a
Map1lc3b
shCtrl
shCtrl-Fluva
shFAM#1
shFAM#1-Fluva
shFAM#2
shFAM#2-Fluva
C
TA
shCtrl
shFAM#1
shFAM#2
Fluva
Mouse ID
FAM134B
GFP
LC3BI
LC3BII
HSP90
kDa
D
TA
Relative protein expression
FAM134B-S/HSP90
GFP/HSP90
LC3B-II/LC3B-I
E
TA
Relative mRNA level
Ccpg1
Atl3
Tex264
ER-phagy receptors
F
Body weight (g)
(Week)
G
Grip strength (% of body weight)
Week1
Week2
Week3
Week4
Week5
Week6
Daily statin treatment
H
Muscle weight (% of body weight)
TA muscle
I
-Fluva
+Fluva
shCtrl
shFAM#1
shFAM#2
Quad
TA
Sol
Gas
J
TA
shCtrl
shFAM#1
shFAM#2
-Fluva
+Fluva
K
Relative frequency (%)
Fiber cross-sectional area (μm²)
L
Cross-sectional area (μm²)
M
TA
SERCA1
SERCA2
RyR
KTN1
Climp63
HSP90
kDa
N
TA
Relative protein expression
SERCA1/HSP90
SERCA2/HSP90
RyR/HSP90
KTN1/HSP90
Climp63/HSP90
O
TA
Relative mRNA level
Serca1
Serca2
Ryr1
SR proteins

**Figure S7. AAV-mediated FAM134B knockdown induces muscle atrophy.**

(A) Schematic of AAV-mediated FAM134B knockdown in the skeletal muscle of mice. The successful injection could be confirmed with GFP signals in the hind limb using a handheld blue LED light with an orange emission filter.

(B) mRNA levels of *Fam134b-L*, *Fam134b-S*, *Map1lc3a* and *Map1lc3b* in the TA. n=5 per group.

(C, D) Immunoblotting images (C) and quantification data (D) of FAM134B, GFP, and LC3B in the TA. HSP90 was used as an internal control. n=4 per group for quantification.

(E) mRNA expression of other ER-phagy receptors, including *Ccpg1*, *Atl3,* and *Tex264* in the TA. n=5.

(F) Body weight was measured weekly during the treatments. n=5.

(G) The grip strength was recorded weekly and normalized to body weight. n=5.

(H) TA weights normalized to body weight. n=5.

(I) Representative images of Quad, TA, Sol, and Gas.

(J-L) H&E staining images (J), distribution of muscle fiber CSA (K), and average muscle fiber CSA (L) in TA are shown (n=290–1,000 fibers from 3 mice each). Scale bar, 50 µm.

(M, N) Immunoblot images (M) and quantification data (N) of SERCA1, SERCA2, RyR, KTN1, and Climp63 in the TA. HSP90 was used as an internal control. n=3 per group.

(O) mRNA expression of *Serca1*, *Serca2*, and *Ryr* in the TA. n=5 per group.

Data shown as mean ± S.D. Statistical analyses were performed by one-way ANOVA with a Tukey-Kramer post hoc test. *, $p < 0.05$, **, $p < 0.01$, ****, $p < 0.0001$.

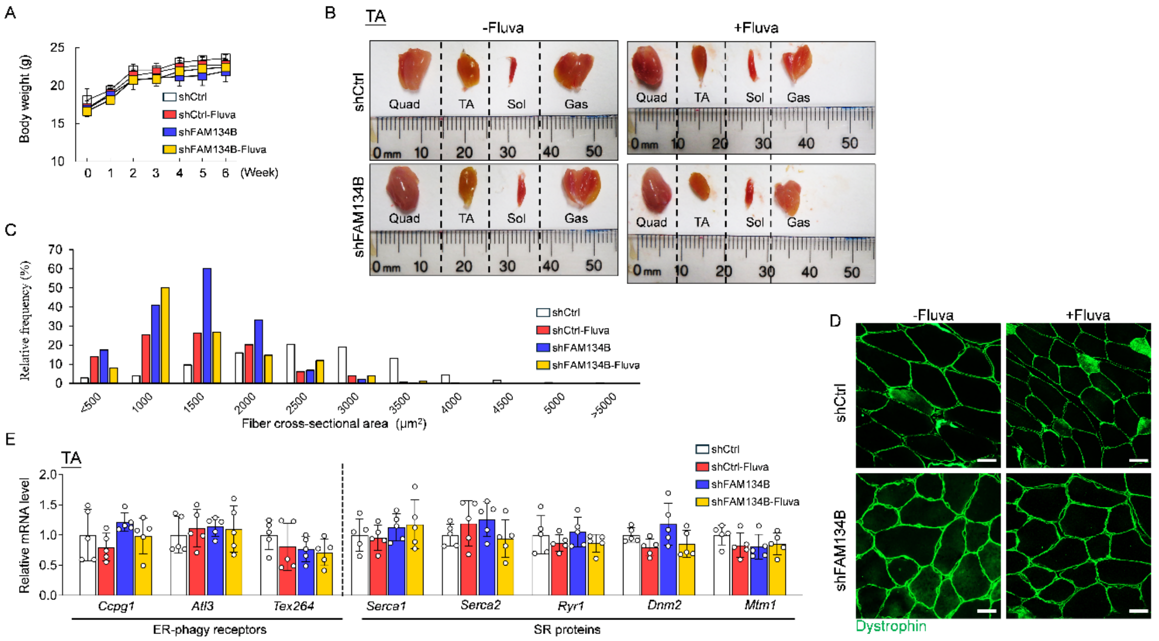


**Figure S8. FAM134B is essential for maintaining the integrity of skeletal muscle.**

(A) Body weight was measured weekly during the treatments. n=5.

(B) Representative images of Quad, TA, Sol, and Gas.

(C) Distribution of muscle fiber CSA in TA, n=485–1101 myofibers from 3 mice each.

(D) Representative images of dystrophin-positive muscle fibers in TA. Scale bar, 20 μm.

(E) mRNA levels of ER-phagy receptors and those for SR/ER proteins in the TA. n=5 per group.

Data shown as mean ± S.D. Statistical analyses were performed by one-way ANOVA with Tukey-Kramer post hoc test. No statistical significance was detected in panels A and E.
